# Repeated-ischaemic exercise enhances mitochondrial and ion transport gene adaptations in human skeletal muscle – Role of muscle redox state and AMPK

**DOI:** 10.1101/156505

**Authors:** Danny Christiansen, Robyn M. Murphy, Jens Bangsbo, Christos G. Stathis, David J. Bishop

**Affiliations:** Institute of Sport, Exercise and Active Living (ISEAL), Victoria University, Melbourne, Victoria, 3011, Australia.; Department of Biochemistry and Genetics, La Trobe Institute for Molecular Science, La Trobe University, Melbourne, Victoria, 3086, Australia.; Department of Nutrition, Exercise and Sports (NEXS), University of Copenhagen, 2200 Copenhagen N, Denmark.; School of Medical and Health Sciences, Edith Cowan University, Perth, Australia

**Keywords:** Ischaemic exercise, PGC-1α, Na^+^-K^+^-ATPase

## Abstract

This study assessed the effect of repeated-ischaemic exercise on the mRNA content of PGC-1α (total, 1α1, and 1α4) and Na^+^,K^+^-ATPase (NKA; α_1-3_, β_1-3_, and FXYD1) isoforms in human skeletal muscle, and studied some of the potential molecular mechanisms involved. Eight trained men (26 ± 5 y and 57.4 ± 6.3 mL·kg^-1^·min^-1^) completed three interval running sessions with (ISC) or without ischaemia (CON), or in hypoxia (HYP, ~3250 m), in a randomised, crossover fashion separated by 1 week. A muscle sample was collected from the dominant leg before (Pre) and after exercise (+0h, +3h) in all sessions to measure the mRNA content of PGC-1α and NKA isoforms, oxidative stress markers (i.e. *catalase* and *HSP70* mRNA), muscle lactate, and phosphorylation of AMPK, ACC, CaMKII, and PLB protein in type I and II fibres. Muscle hypoxia (i.e. deoxygenated haemoglobin) was matched between ISC and HYP, which was higher than in CON (~90% vs. ~70%; p< 0.05). The levels of *PGC-1α* total, *-1α1, −1α4*, and *FXYD1* mRNA increased in ISC only (p< 0.05). These changes were associated with increases in oxidative stress markers and higher p-ACCSer^221^/ACC in type I fibres, but were unrelated to muscle hypoxia, lactate, and CaMKII and PLB phosphorylation. These findings highlight that repeated-ischaemic exercise augments the skeletal muscle gene response related to mitochondrial biogenesis and ion transport in trained men. This effect seems attributable, in part, to increased oxidative stress and AMPK activation, whereas it appears unrelated to altered CaMKII signalling, and the muscle hypoxia and lactate accumulation induced by ischaemia.

**Summary in key points:**

- We investigated if ischaemia would augment the exercise-induced mRNA response of PGC-1α and Na^+^,K^+^-ATPase (NKA) isoforms (α_1-3_, β_1-3_, and FXYD1), and examined whether this effect could be related to oxidative stress and fibre type-dependent AMPK and CaMKII signalling in the skeletal muscle of trained men.
- Repeated-ischaemic exercise increased the mRNA content of PGC-1α total, −1α1, and-1α4, and of the NKA regulatory subunit FXYD1, whereas exercise in systemic hypoxia or alone was without effect on these genes.
- These responses to ischaemia were complemented by increased oxidative stress (as assessed by *catalase* and *HSP70* mRNA) and ACC phosphorylation (an indicator of AMPK activation) in type I fibres. However, they were unrelated to CaMKII signalling, muscle hypoxia, and lactate accumulation.
- Thus, repeated ischaemic exercise augments the muscle gene response associated with mitochondrial biogenesis and ion homeostasis in trained men. This effect seems partly attributable to promoted oxidative stress and AMPK activation.

**Abbreviations:**

ACC: Acetyl-CoA carboxylase
AMPK: 5’ AMP-activated protein kinase subunit
β2M: β2 microglobulin
CaMKII: Ca2+-calmodulin-dependent protein kinase isoform II
CON: control session
C_T_: cycle threshold
CV: coefficient of variation
FXYD1: phospholemman isoform 1
GAPDH: glyceraldehyde 3-phosphate dehydrogenase
GXT: graded exercise test
HHb: deoxygenated haemoglobin
HSP70: heat-shock protein 70
HYP: repeated-hypoxic exercise session
ISC: repeated-ischaemic exercise session
K^+^: potassium ion
LT: lactate threshold
MHC: myosin heavy chain
Na^+^: sodium ion
NIRS: near-infrared spectroscopy
NKA: Na^+^, K^+^-ATPase
OXPHOS: oxidative phosphorylation
PGC-1α: peroxisome proliferator-activated receptor-gamma coactivator 1 alpha
PLB: phospholamban
ROS: reactive oxygen species
SDS: sodium dodecyl sulphate
TBP: TATA-binding protein
VO_2max_: maximum oxygen uptake

## Introduction

The movement pattern of many sports (e.g., football codes, handball, and tennis) requires athletes to possess a well-developed ability to repeatedly perform maximal or near-maximal intensity efforts (Povoas *et al.*, 2012; Varley *et al.*, 2013; Adriano Pereira *et al.*, 2016). This ability appears limited by factors intrinsic to skeletal muscle (Bangsbo, 1994; Bishop, 2012; Bishop & Girard, 2013). For example, fatigue during this type of activity has been related to perturbations in muscle transmembrane sodium (Na^+^) and potassium ion (K^+^) gradients, which exceed the transport capacity of the Na^+^,K^+^-ATPase (NKA) to sustain transmembrane ion equilibrium (McKenna, 1992; McKenna *et al.*, 2008). As such, the NKA is important for performance during intense intermittent exercise. The NKA is composed of a catalytic α, a structural β, and a regulatory γ (phospholemman or FXYD) subunit (Clausen, 2013), all of which are expressed as different isoforms (α_1-3_, β_1-3_ and FXYD1) in human skeletal muscle (Wyckelsma *et al.*, 2015). The relative distribution of these isoforms determine, in part, the potential to generate active NKA complexes at the cell surface. Therefore, changes in the expression of different NKA isoforms could affect the muscle’s capacity for Na^+^/K^+^ transport (Nielsen *et al.*, 2004).

Another limiting factor for the ability to perform repeated, intense exercise is the muscle’s capacity to generate ATP via oxidative pathways (Thomas *et al.*, 2004). Accordingly, increases with exercise training in both mitochondrial respiratory function and content (as assessed by citrate synthase activity) have been temporally associated with an enhanced exercise capacity (Bishop *et al.*, 2014). A key determinant of these endurance-type adaptations is the transcriptional co-activator, the peroxisome proliferator-activated receptor-γ coactivator 1α (PGC-1α). It has recently been reported in humans that one isoform of PGC-1α, namely the isoform 1 (PGC-1α1), orchestrates exercise-induced increases in the content of mitochondrial genes (e.g. *OXPHOS*), whereas another isoform, the isoform 4 (PGC-1α4), has been shown to regulate the expression of genes that influence muscle hypertrophy (Ruas *et al.*, 2012). The expression of specific isoforms of PGC-1α could determine the nature of the responsiveness of skeletal muscle to various external stimuli, such as exercise.

Despite a scarcity of published research, it is believed that repeated, transient bursts in gene expression enhances the potential for increasing the protein abundance of NKA and PGC-1α isoforms in human skeletal muscle (Perry *et al.*, 2010; Christiansen et al., 2017, unpublished). Identifying the cellular stressors involved in the transcription of these isoforms would therefore seem important. However, at present, the cellular stressors essential for the upregulation of NKA and PGC-1α isoforms in human muscle are poorly defined. What is known, primarily based on cell culture and animal models *in vitro*, is that the transcription of these isoforms might be initiated by ionic and metabolic perturbations, including shifts in K^+^ (Zhuang *et al.*, 2000; Wang *et al.*, 2007), lactate (Hashimoto *et al.*, 2007), and Ca^2+^concentrations (Ojuka *et al.*, 2003; Norrbom *et al.*, 2004), and by the formation of reactive oxygen species (ROS) (Wendt *et al.*, 1998; Murphy *et al.*, 2008; Irrcher *et al.*, 2009). These cellular stressors have been shown to activate several signalling kinases, of which the AMP-stimulated protein kinase (AMPK), the Ca^2+^calmodulin-dependent protein kinase (CaMK), or both, have been coupled to the transcriptional regulation of PGC-1α (Wu *et al.*, 2002; Irrcher *et al.*, 2008) and NKA isoforms (Nordsborg *et al.*, 2010a). Thus, strategies promoting perturbations in ionic gradients, metabolic and redox homeostasis, and ROS production, could prove useful for increasing the muscle’s expression of NKA- and PGC-1α-isoform mRNA transcripts.

One potential strategy to augment these cellular stressors might be to exercise with reduced muscle blood flow (ischaemia), known as ischaemic exercise. Ischaemia has typically been invoked by inflation of an occlusion cuff fixed around the limb(s) proximal to locomotor muscles and has been applied during multiple exercise modes, including walking, cycling and resistance training (Abe *et al.*, 2006; Abe *et al.*, 2010; Cook *et al.*, 2014). Inflation of the cuff has been reported to compromise both the arterial and venous flow (Sundberg & Kaijser, 1992; Horiuchi & Okita, 2012), resulting in an hypoxic and more acidic intramuscular environment (Sundberg & Kaijser, 1992). A successive deflation of the cuff can promote local reactive hyperaemia (Gundermann *et al.*, 2012). In combination, these mechanisms seem a powerful stimulus for amplifying the transient bursts in skeletal muscle ROS production that accompany consecutive bouts of exercise (Slezak *et al.*, 1995; Clanton, 2007). They could also affect K^+^ and calcium ion (Ca^2+^) concentrations by modulating the function of ion channels and transport systems (Kourie, 1998; Juel *et al.*, 2015). There is therefore evidence to suggest ischaemic exercise could be an effective strategy to promote the mRNA expression of NKA and PGC-1α isoforms in human skeletal muscle. However, this hypothesis remain presently untested.

The primary aim of the present study was therefore to explore the effect of ischaemic exercise on the mRNA content of PGC-1α (total, and isoforms 1α1 and 1α4) and NKA isoforms (α_1-3_, β_1-3_, and FXYD1) in human skeletal muscle. Our second aim was to elucidate some of the potential molecular mechanisms involved. Our working hypotheses were: 1) Ischaemia would augment the effect of exercise on PGC-1α- and NKA-isoform expression, and 2) increases in the expression of these isoforms would coincide with elevated levels of oxidative stress markers (i.e. *catalase* and *HSP70* mRNA content) and Acetyl-CoA carboxylase (ACC; an indicator of AMPK activation) and CaMKII phosphorylation. Evidence from astrocytes *in vitro* suggests that the reperfusion phase, rather than hypoxia *per se*, may be the primary stimulus underlying increases in the mRNA content of NKA isoforms in response to hypoxia-reperfusion (Kasai *et al.*, 2003). Thus, we also included a hypoxic condition (i.e. exercising in normobaric, systemic hypoxia) to assess the hypothesis that 3) increases in isoform expression could not be attributed to the muscle hypoxia induced by occlusion.

## Methods

### Ethical Approval

This study was approved by the Human Research Ethics Committee of Victoria University, Melbourne, Australia (HRE14-309), and was performed in accordance with the latest instructions in the *Declaration of Helsinki*. Participants provided oral and written informed consent before enrolment in the study.

### Participants

Eight healthy men engaged in team sports (five in soccer, two in Australian-rules football, and one in basketball) participated in the study. Their physical characteristics are shown in Table 1. All participants were non-smokers and engaged in their sport 3-5 times per week.

**Table 1:**
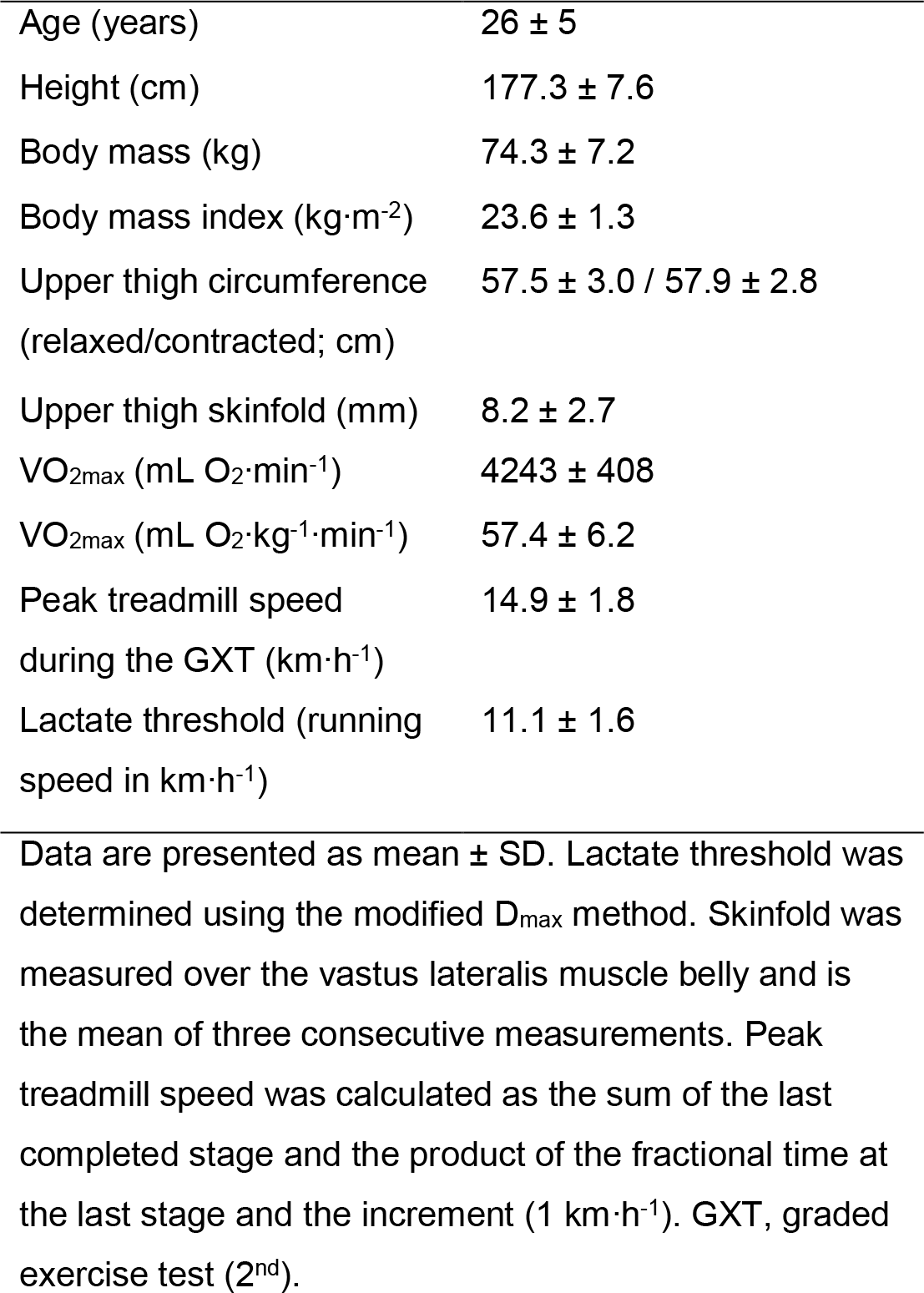
Participant characteristics

### Randomisation and blinding

The study was a randomised, counterbalanced, crossover experiment, and took place in the Exercise Physiology Laboratory at the Institute of Sport, Exercise and Active Living (ISEAL), Victoria University, Melbourne, Australia. All sessions were performed on a Katana Sport XL treadmill (Lode, Groningen, Netherlands) in 21.4 ± 1.1 °C and 40.8 ± 6.8 % humidity. Participants completed three main trials matched for total work, duration (34 min), and work:rest ratio. These trials were separated by one week and consisted of interval running with (ISC) or without ischaemia (CON), or in normobaric, systemic hypoxia (HYP). Each participant was allocated a trial order using a random-number generator (MS Excel 2013, Microsoft, USA). To minimise any perceived placebo effect (not to be confused with a true placebo effect) (Ernst & Resch, 1995), the participants were not informed about which trial was hypothesised to be of greatest value to the physiological response, and whether they were breathing hypoxic or normoxic air. A pneumatic tourniquet (Riester, Germany) was attached to the participants’ preferred kicking (dominant) leg by adhesive tape in all trials, but it was only inflated in ISC. In addition, the participants were informed that the study purpose was to evaluate the effect of different degrees of ischaemia. Information about what trial was to be performed on each occasion was given on the day of execution.

### Pre-testing

Prior to the main trials, the participants visited the laboratory on four separate occasions interspersed by at least 48 h. On the first and fourth visit, participants performed a graded exercise test (GXT). This test was used to assess the participants’ lactate threshold (LT) and maximum oxygen consumption (VO_2max_). The LT from the fourth visit was used to determine individual running speed during the main trials (i.e. ISC, CON and HYP). On the second visit, participants performed the ISC trial with near-infrared spectroscopy (NIRS) probes placed over the vastus lateralis muscle belly of their dominant leg to assess muscle oxygen content (cf. section on *Muscle deoxygenation*), and to accustom the participants to ISC and the equipment. During the third visit, participants completed the same running protocol with NIRS probes attached. The first three exercise bouts during this visit were performed without ischaemia or hypoxia. The remaining six bouts were completed in normobaric, systemic hypoxia to accustom the participants to HYP and to allow estimation of individual inspired oxygen fraction (FiO2) to be used in HYP to match the level of muscle hypoxia during ISC (detailed in *Ischaemia and hypoxia*). The tourniquet was worn during both the second and third visit.

### Main trials

On the days of the main trials, the participants reported to the laboratory between 8-9 am after 7.3 ± 1.1 h of sleep and after consuming a standardised dinner and breakfast (detailed in *Diet and activity control*) 15 h and 2.5 h, respectively, prior to arrival. After approximately 30 min of rest in the supine position, a catheter was inserted in an antecubital vein, allowing mixed-venous blood to be sampled. After an additional 15 min of rest, blood and muscle was sampled, also in the supine position. Next, the participants moved to the treadmill where they were instrumented with one pair of NIRS optodes on the belly of the vastus lateralis muscle of their preferred kicking (dominant) leg to reliably and non-invasively monitor muscle deoxygenation *in vivo* (Van Beekvelt *et al.*, 2001). A belt was placed around their chest to measure heart rate. In a sitting position with the dominant leg unloaded, muscle deoxygenation was measured for at least 2 min until a plateau was reached and a stable baseline reading was recorded. Next, participants were fitted with a facemask covering the mouth and nose to enable them to breathe normoxic or hypoxic air. A pneumatic tourniquet was attached to the participants’ dominant leg by adhesive tape. The tourniquet was inflated only in ISC before each bout of exercise and deflated upon termination of each bout. A time-aligned schematic representation of the experimental protocol is shown in Fig. 1. Every trial commenced with a 5-min warm-up (WU) at 75 % LT followed by 5 min of rest. At the third and fourth minute of the WU, a 5-s acceleration to ~110 % LT, followed by a 5-s deceleration to 75 % LT, was performed. Next, three series of three 2-min runs were executed at a fixed relative intensity (105 % LT, 11.6 ± 1.7 km·h^-1^; no incline). The runs were separated by 1 min, and the series by 5 min, of walking (~5 km·h^-1^), respectively. This design was introduced to promote repeated, transient bursts in ROS formation (Slezak *et al.*, 1995; Raedschelders *et al.*, 2012), which is a key stimulus for increases in the mRNA content of NKA and PGC-1α isoforms in cell culture (Silva & Soares-da-Silva, 2007; Irrcher *et al.*, 2009). The duration of runs and the work:rest ratio were based on pilot work and balanced to achieve the highest tolerable mean speed, by which the exercise regimen could be completed with the chosen magnitude of leg ischaemia (3.0 mmHg·cm^-1^). This training regimen (2:1 min) has been demonstrated to increase Na^+^-K^+^-ATPase content (as assessed by [^3^H]-ouabain binding) and enhance intense intermittent exercise performance (Edge *et al.*, 2013). Antecubital venous blood was sampled at rest before exercise, prior to each series, immediately after each run, and at 3 and 6 min after termination of exercise. Muscle was sampled at rest in the supine position before (Pre), immediately after (64 ± 28 s; +0h), and 3 h post, exercise (+3h).

**Figure 1.**
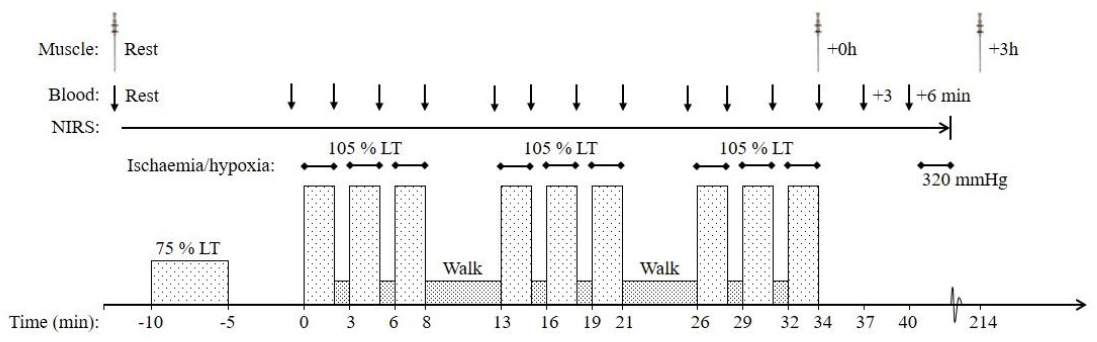
Time-aligned, schematic representation of the experimental protocol. The participants performed three exercise trials separated by one week consisting of running without (control) or with the muscle blood flow partially occluded (ischaemia), or in normobaric, systemic hypoxia (hypoxia). The intensity of exercise was set according to the participants’ individual lactate threshold (LT). Muscle was sampled at rest before, immediately post, and after 3 h of recovery from, each trial. Blood was sampled from an antecubital vein at the time points indicated.

### Ischaemia and hypoxia

In all trials, a pneumatic tourniquet made of nylon with a width of 13 cm (Riester, Germany) was externally applied to the most proximal part of the participants’ preferred kicking leg. In ISC, 15 s prior to the onset of a run, the tourniquet was rapidly inflated over ~10 s to reach an end-pressure of 3.0 mmHg·cm^-1^ (i.e. relative to thigh circumference, TCF). The mean pressure was in the lower end of the range of pressures used in previous studies (~3-5 mmHg·cm^1^) (Abe *et al.*, 2005; Abe *et al.*, 2006; Fujita *et al.*, 2008; Abe *et al.*, 2010; Kubota *et al.*, 2011). The pressure during running ranged from (mean ± SD) 123 ± 12 (range: 109-139) mmHg in the float phase to 226 ± 24 (range: 200-260) mmHg in the landing phase. The difference between our predetermined (~175 mmHg) and actual (mean ± SD) pressure during the trials was −1 ± 8.5 mmHg. The tourniquet was deflated immediately after termination of exercise. After 15 min of recovery from exercise, the tourniquet was inflated to 320 mmHg until a maximum plateau in muscle deoxygenation. TCF was measured before exercise as one third of the distance midline from the inguinal crease to the proximal boarder of patella. This represented the site of tourniquet application. In HYP, the participants executed the exercise bouts in normobaric, systemic hypoxia with an F_i_O_2_ of 14.0 %, corresponding to an altitude of approximately 3250 m.

### Muscle deoxygenation

Deoxygenation at the muscle level was measured by continuous-wave, near-infrared spectroscopy (NIRS), as described previously (Van Beekvelt *et al.*, 2001). A pair of NIRS optodes was positioned over the distal part of the vastus lateralis muscle ~15 cm above the proximal boarder of patella. Optodes were fixed in a plastic spacer, which was attached to the skin by double-sided sticky disks to ensure direct contact between optodes and skin. A black bandage was placed over the optodes and around the leg for further fixation and to shield against extraneous light, and to minimise loss of transmitted near-infrared light. The interoptode distance was 40 mm. Skinfold thickness was measured between the emitter and receiver optodes using a skinfold caliper (Harpenden Ltd.). Skinfold thickness (8.2 ± 2.7 mm) was less than half the distance separating the optodes. Circumference of the plastic spacer was marked on the skin using an indelible pen and pictures were taken to ensure that optodes were placed at the same position in all trials. Light absorption signals were converted to HHb deoxygenation changes using a differential pathlength factor (DPF) calculated according to participants’ age. The DPF was the same across trials for each participant. Data were acquired at 10 Hz and subsequently filtered in R software (ver. ×64 3.2.5, R Foundation for Statistical Computing) using a 10^th^ order zero-lag, low-pass Butterworth filter with a cut-off frequency of 0.1. The optimal cut-off frequency (i.e. reducing over- and underestimation of local means) was predetermined by an iterative analysis of root-mean-squared residuals derived from the application of multiple filters by use of a range of cut-off frequencies (0.075-0.150). Filtered data was used for the final analysis. Time-alignment and normalisation to the signal range between baseline (resting) and maximum (full occlusion) readings were completed in Excel (Ver. 2013, MS Office, Microsoft, USA).

### Graded exercise test (GXT)

Participants completed the GXT following a light standardised meal ~3 hours prior to arrival. The test consisted of 4-min runs punctuated by 1 min of rest. The first run commenced at 5.0 km·h^-1^, and the second at 8 km·h^-1^ The speed was then increased by 1 km·h^-1^ at the onset of each subsequent run until volitional exhaustion, defined as an inability to maintain the required speed. This progression in speed allowed a minimum of seven running stages to be completed (range: 7-11). After 5 min of rest, participants commenced running at the speed of the last completed run, after which the speed was increased by 1 km·h^-1^ per minute until volitional exhaustion. This incremental bout was performed to ascertain attainment of a maximum 30-s plateau in oxygen consumption. Before the test, a facemask was placed over the mouth and nose and connected to an online, gas-analysing system for measurement of inspired and expired gases. To determine LT, blood was sampled at rest and immediately after each running stage from a 20-gauge, antecubital venous catheter. The catheter was inserted at rest in a supine position on a laboratory bed at least 20 min prior to the test. The LT was calculated using the modified D_max_ method as it has been shown to better discriminate between individuals in comparison with other methods (Bishop *et al.*, 1998). VO_2max_ was determined as the mean of the two peak consecutive 15-s values recorded during the test.

### Diet and activity controls

Participants consumed a standardised dinner (55 kJ·kg^-1^ BM; 2.1 g carbohydrate·kg^-1^ BM, 0.3 g fat·kg^-1^ BM, and 0.6 g protein·kg^-1^BM) and breakfast (41 kJ·kg^-1^ BM; 1.8 g carbohydrate·kg^-1^ BM, 0.2 g fat·kg^-1^BM, and 0.3 g protein·kg^-1^ BM) 15 and 3 hours, respectively, before every main trial. They recorded their dietary pattern within 48 h prior to each laboratory visit and were asked to replicate the same nutritional intake as per before their first exercise trial. Participants were instructed to maintain their normal dietary pattern throughout the study and were free of anti-inflammatory drugs and supplements, as well as medicine. The participants were instructed to replicate their weekly routine physical activity throughout the study and to avoid activity beyond daily living in the 48 h prior to each visit. In the 3-h period from termination of exercise to the +3h biopsy, oral consumption was limited to *ad libitum* water.

### Muscle sampling

Vastus lateralis muscle biopsies were collected from the dominant leg in all trials using the Bergström needle biopsy technique with suction, amounting to 9 biopsies per participant. To minimise bleeding, the biopsy in ISC was obtained immediately after deflation of the tourniquet. In preparation for a muscle sample, a small incision was made under local anaesthesia (5 ml, 1% Xylocaine) through the skin, subcutaneous tissue and fascia of the muscle. Incisions were separated by approximately 1-2 cm. Immediately after sampling, samples were rapidly blotted on filter paper to remove excessive blood, and instantly frozen in liquid nitrogen. The samples were stored at −80° C until being analysed. The incisions were covered with sterile Band-Aid strips and a waterproof Tegaderm film dressing (3M, North Ryde, Australia).

### Blood handling and analysis

To ensure blood samples accurately represented circulating blood, ~2 mL of blood was withdrawn and discarded before sampling of approximately 2 mL of blood per sample. After being drawn, samples were instantaneously placed on ice until being analysed for lactate, pH, and K^+^, concentrations after exercise on an ABL 800 Flex blood gas analyzer (Radiometer, Brønshø j, Denmark).

### Arterial oxygen saturation (S_a_O_2_)

For safety reasons, adhesive optodes were placed on the tip of the left index finger to monitor arterial oxygen saturation during the HYP trial by pulse oximetry (Nellcor N-600, Nellcor Inc., Hayward, CA). Data was recorded at rest in the standing position on the treadmill, and during the final minute of each bout of running.

### RNA isolation and reverse transcription

Approximately 44 mg w.w. muscle per sample was homogenised (2 × 2 min at 30 Hz) in ~800 μ L TRIzol reagent (Invitrogen, Carlsbad, CA) using an electronic homogeniser (FastPrep FP120 Homogenizer, Thermo Savant). After homogenisation, the supernatant was aspirated into a new, freshly autoclaved microfuge tube containing 250 μL chloroform (Sigma Aldrich, St. Louis, MO). After few manual inversions and 5 min on ice, the mixture was centrifuged (15 min at 13.000 rpm) at 4°;C. After centrifugation, the superior phase was pipetted into a new, autoclaved microfuge tube, and 400 μ L 2-isopropanol alcohol (Sigma-Aldrich, St Louis, MO) and 10 μ L of 5 M NaCl were added. The samples were then stored at −20°C for 3 h to precipitate the amount of RNA. After cooling, the samples were centrifuged (20 min at 13.000 rpm) at 4°C, and the isopropanol aspirated. The RNA pellet was rinsed with 75 % ethanol made from DEPC-treated H_2_O (Invitrogen Life Sciences), and centrifuged (8 min at 9000 rpm) at 4°C. After pipetting off the ethanol, the pellet was resuspended in 5 μ L of heated (60 °C) DEPC-treated H_2_O. The samples were stored at −80°C until reverse transcription. RNA purity (mean ± SD, 1.96 ± 0.24; 260nm/280nm) and concentration (mean ± SD, 1317.4 ± 1311.5 ng·μL^-1^) were determined spectrophotometrically on a NanoDrop 2000 (Thermo Fisher Scientific, Wilmington, DE). In addition, RNA integrity was assessed in six randomly chosen samples on an electrophoresis station (Experion, BioRad) using the manufacturer’s RNA analysis kit (Experion RNA StdSens) and instructions. The RNA quality indicator (RQI) of the six samples was (mean ± SD) 8.1 ± 0.7. One microgram of RNA per sample was reverse-transcribed into cDNA on a thermal cycler (S1000™ Thermal Cycler, Bio-Rad, Hercules, CA) using a cDNA synthesis kit (iScript RT Supermix, #1708841; Bio-Rad, Hercules, CA). The following incubation profile was used with random hexamers and oligo dTs in accordance with the manufacturer’s instructions: 5 min at 25 °C, 20 min at 46 °C and 1 min at 95 °C. cDNA was stored at −20°C until real-time PCR.

### Real-time RT-PCR

Real-time PCR was performed to determine the expression of target and reference genes. Reactions were prepared on a 384-well plate using an automated pipetting system (epMotion 5073l, Eppendorf, Hamburg, Germany). One reaction (5 μL) was composed of 2 μL diluted cDNA, 0.15 μL forward and reverse primer (100 μM concentration), 0.2 μL DEPC-treated H_2_O, and 2.5 μL iTaq universal SYBR Green Supermix (#1725125; Bio-Rad, Hercules, CA). Real-time PCR was performed on a QuantStudio 7 Flex Real-Time PCR System (#4485701, Thermo Fisher Scientific, USA) using the following protocol: Denaturing at 95 °C for 3 min, followed by 40 cycles of 95 °C for 15 s, and 60 °C for 60 s. Reactions were run in duplicate on the same plate with four template-free and four RT-negative controls. The expression of target genes was normalised to that of three reference genes using the 2*^-ΔΔC_T_^* method (Livak & Schmittgen, 2001). This correction has been shown to yield reliable and valid mRNA data (Vandesompele *et al.*, 2002). Reference genes used were *glyceraldehyde 3-phosphate dehydrogenase* (*GAPDH*), *TATA-binding protein* (*TBP*), and *β2 microglobulin* (*β2M*). The mean (± SD) coefficient of variation (CV) of duplicate reactions (CT), along with the forward and reverse sequences for the primers are shown in Table 2. Criteria and procedure for the design of primers for NKA isoforms are presented elsewhere (Christiansen *et al.* 2017, in review, *FASEB J*). Primers for PGC-1α isoforms were identical to those previously used (Ruas *et al.*, 2012). Primer specificity was confirmed by performing a melt curve analysis at the end of each PCR run.

**Table 2:**
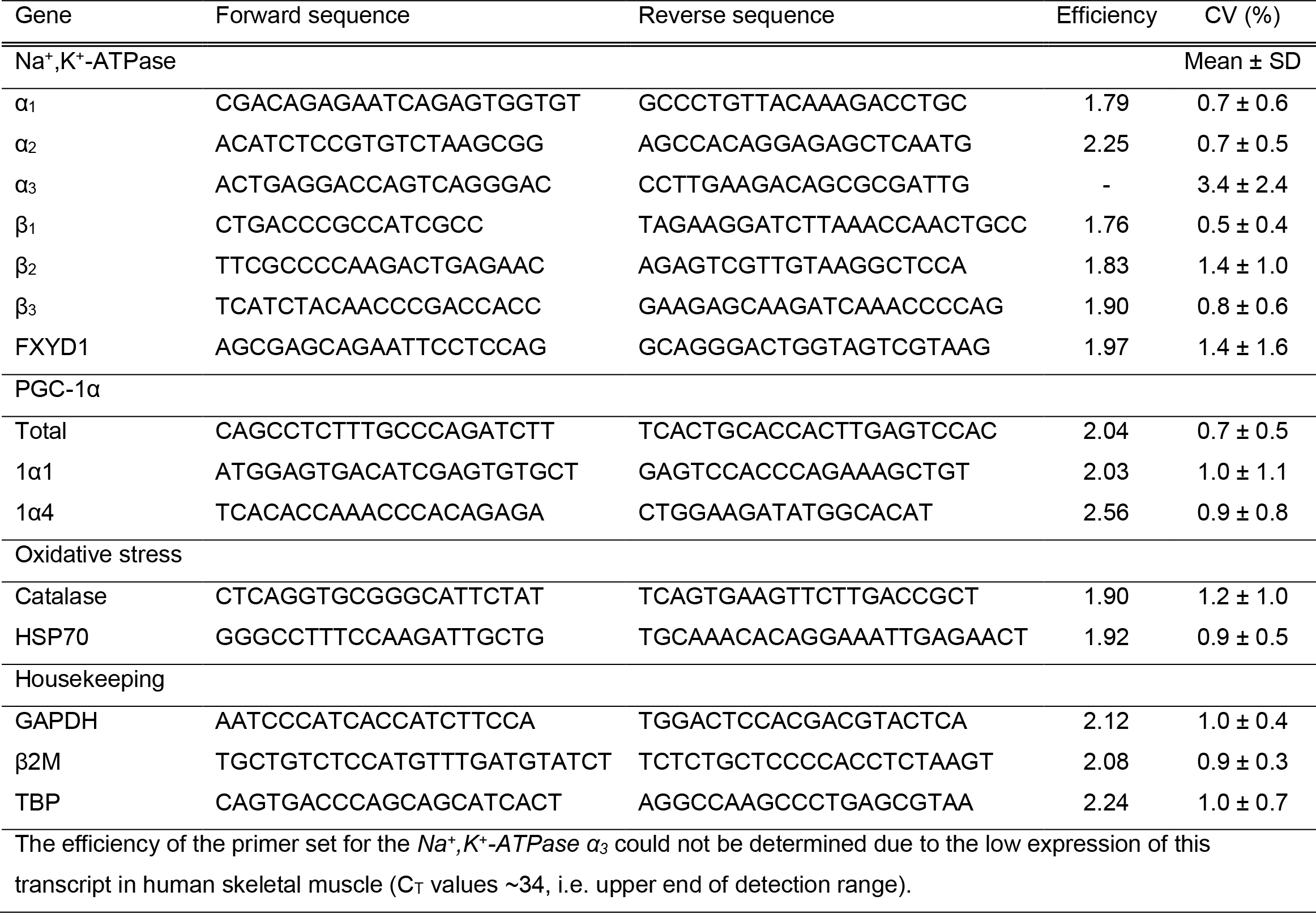
Forward and reverse primer sequences used in real-time PCR, their amplification efficiency, and the coefficient of variation (CV) of duplicates.

### Dissection and fibre typing of muscle fibres

All chemicals used for dot blotting and Western blotting were from Bio-Rad unless otherwise stated. Antibodies are detailed in Table 3.

**Table 3:**
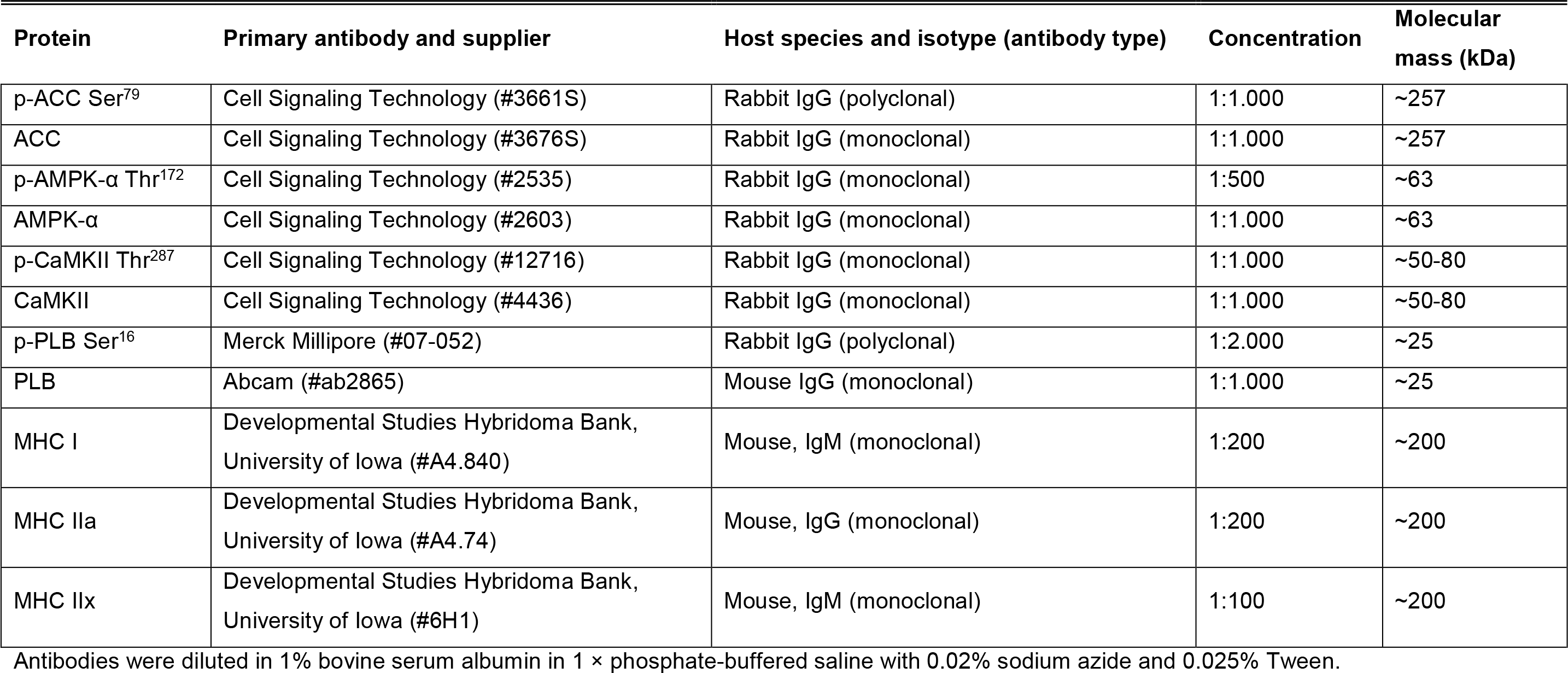
Primary antibodies used for dot blotting and western blotting

One part of each muscle biopsy (49.6 ± 10.0 mg w.w.) was freeze dried for 40 h, yielding 11.6± 2.7 mg d.w. muscle tissue. From these freeze-dried portions, a minimum of 40 single-fibre segments per sample (range: 40-120; total *n* = 2750) were isolated under a dissecting microscope using fine jeweller’s forceps. The segments were placed in individual microfuge tubes and incubated instantly for 1 h at room temperature in 10-μL denaturing buffer (0.125 M Tris-HCl, 10% glycerol, 4% SDS, 4 M urea, 10% mercaptoethanol, and 0.001% bromophenol blue, pH 6.8), in accordance with previous procedure (Murphy, 2011). The denatured segments were stored at −80°C until future use.

Fibre type of individual segments was determined using dot blotting, as recently described (Christiansen et al. 2017, *in preparation*). In brief, two 0.45-μm PVDF membranes were activated in 95 % ethanol and equilibrated for two minutes in cold transfer buffer (25 mM Tris, 192 mM glycine, pH 8.3, 20 % methanol), after which a 1.5 μL aliquot of denatured sample, corresponding to one seventh of a fibre segment, was spotted onto each membrane. The membranes were placed at room temperature on a dry piece of filter paper to dry completely (5-10 min), after which they were reactivated in the ethanol and re-equilibrated in transfer buffer. After a quick wash in Tris-buffered saline-Tween (TBST), membranes were blocked in 5 % non-fat milk in TBST (blocking buffer) for 5-15 min. One of the blocked membranes was incubated (1 in 200 in 1 % BSA with PBST) with MHCI antibody, and the other membrane with MHCIIa antibody for 2 h at room temperature with gentle rocking. After a quick wash in blocking buffer, membranes were incubated (concentration: 1:20.000) with goat anti-mouse IgG (MHCIIa, #PIE31430, ThermoFisher Scientific) or IgM (MHCI, #sc-2064, Santa Cruz Biotechnology, TX, USA) horse radish peroxidase (HRP)-conjugated secondary antibody for 1 h at room temperature with rocking. Membranes were then quickly rinsed in TBST, exposed to Clarity enhanced chemiluminescence reagent (Bio-Rad, Hercules, CA, USA), and imaged on a ChemiDoc MP (Bio-Rad). The membrane incubated with MHCIIa antibody was reprobed with MHCIIx antibody for 2 h with rocking at room temperature, after which it was exposed to the same secondary antibody as MHCI (#sc-2064, Santa Cruz Biotechnology, TX, USA) for 1 h at room temperature and imaged accordingly. The difference in the host immunoglobulin species of the MHCIIa (IgG) and MHCIIx (IgM) antibodies allowed both isoforms to be quantified on the same membrane.

The remnant of each denatured fibre segment (7 μL) was grouped according to MHC expression to form samples of type I (MHCI) and type II (MHCIIa) fibres for each biopsy, in line with previous procedure (Kristensen *et al.*, 2015). The number of fibre segments included in each group of muscle fibres per biopsy was (mean ± SD) n = 12 ± 6 (range: 5-27) for type I, and n = 16 ± 5 (range 7-33) for type IIa, fibres. Hybrid fibres (expressing multiple MHC isoforms) and type IIx fibres (classified by absence of MHCI and MHCIIa, but presence of MHCIIx protein), both constituting 3.1 % of the total pool of fibres, were excluded from analysis.

### Immunoblotting

Fibre-type specific protein abundance and phosphorylation status of the signalling proteins AMPK and CaMKII, and their downstream targets ACC and PLB, respectively, were determined by Western blotting. Fifteen micrograms of protein per sample (~5 μL) were separated (45 min at 200 V) on 26-wells, 4-15 % Criterion TGX stain-free gels (Bio-Rad, Hercules, CA). Each gel was loaded with all samples from one participant, two calibration curves (a four- and a three-point to improve the reliability) and two protein ladders (PageRuler, Thermo Fischer Scientific). Calibration curves were of human whole-muscle crude homogenate with a known protein concentration, which was predetermined as described previously (Christiansen *et al*. 2017, in review, *The Journal of Physiology*). After electrophoresis, gels were UV activated (5 min) on a Criterion stain-free imager (Bio-Rad). Proteins were wet-transferred to 0.45 μm nitrocellulose membrane (30 min at 100 V) in a circulating bath at 4 °C in transfer buffer (25 mM Tris, 190 mM glycine and 20 % methanol). Membranes were then incubated (10 min) in antibody extender solution (Pierce Miser, Pierce, Rockford, IL, USA), washed in double-distilled H_2_O, and blocked for 2 h in blocking buffer (5 % non-fat milk in Tris-buffered saline Tween, TBST) at room temperature with rocking. To allow multiple proteins to be quantified on the same membrane, the membranes were cut horizontally at appropriate molecular masses using the two protein ladders as markers prior to probing with the primary antibodies overnight at 4 °C, and for 2 h at room temperature with constant, gentle rocking. Antibody details are presented in Table 3. Primary antibodies were diluted in 1 % bovine serum albumin (BSA) in phosphate-buffered saline with 0.025% Tween (PBST) and 0.02% NaN_3_. After washing in TBST and probing with appropriate horseradish peroxidase (HRP)-conjugated secondary antibody (goat anti-mouse immunoglobulins or goat anti-rabbit immunoglobulins; Pierce, Rockford, IL, USA) for 1 h with rocking at room temperature, chemiluminescent images of membranes were captured on a ChemiDoc Touch (Bio-Rad), followed by densitometry using Image Lab (Ver. 5.2.1, Bio-Rad). Protein ladders were captured under white light prior to chemiluminescent imaging without moving the membranes. Band densities for proteins were quantified by reference to the linear calibration curves on the same gel and normalised to the total amount of protein in each lane on the stain-free gel image. The same researcher with substantial experience with the techniques was responsible for performing all muscle analyses.

### Muscle ATP and lactate

A portion of each freeze-dried muscle sample (2 mg d.w.) was dissected free of connective tissue, blood and fat before being powdered using a Teflon pestle. The content of ATP and lactate in each sample was extracted using precooled perchloric acid/EDTA and KHCO_3_, and analysed fluorometrically using a modification of a method previously described (Harris *et al.*, 1974), where samples are analysed in a 96-well plate format. All samples from each participant, along with two standards of either ATP or lactate, a four-point NADH standard curve, and blanks (i.e. double distilled H2O) were analysed in triplicate on the same plate. Absorbance readings of samples were normalised to the standards and subtracted blanks.

### Statistics

Data was firstly assessed for normality using the Shapiro-Wilk test. An appropriate transformation of data was applied, if necessary, to obtain a normal distribution prior to subsequent statistical analyses. A two-way repeated-measures analysis of variance (RM ANOVA) was used to test the null-hypothesis of no effect of time (Pre, +3h) and condition (CON, ISC, HYP) for mRNA data (using the 2^-ΔΔCt^ expression), and for blood and metabolite data. The sample size used for gene analyses was n = 8 for the CON. Due to contamination (and a C_T_ >35) of some samples, the sample size was reduced to n = 6 and n = 5 for the ISC and HYP, respectively. A one-way RM ANOVA was used to test the null-hypothesis of no effect of condition for mRNA changes with time using ΔmRNA values (i.e. difference between Pre and +3h). A two-way RM ANOVA was used to test the null-hypothesis of no effect of time (Pre, +0h) and fibre type (type I and type II) within condition for the content and phosphorylation of proteins and to evaluate conditional interactions with time (Pre, +0h) within fibre type. Data normalised to total protein, and not relative changes, was used for protein analyses. For NIRS data, a two-way RM ANOVA was used to test the null-hypothesis of no time and condition effects using the Butterworth-filtered data. Multiple pairwise, *post hoc* analyses used the Tukey test. Interpretation of effect size (*d*) was based on Cohen’s conventions, where < 0.2, 0.2-0.5, >0.5-0.8 and >0.8 were considered as trivial, small, moderate and large effect, respectively (Cohen, 1988). Data are reported as means ± SEM unless otherwise stated. The α-level was set at p ≤ 0.05. Statistical analyses were performed in Sigma Plot (Ver. 11.0, Systat Software, CA).

## Results

### PGC-1α gene transcripts

*PGC-1α* total mRNA increased from Pre to +3h in ISC (p = 0.007, d = 1.32), but was not significantly altered in CON (p =0.447, d = 1.06) or in HYP (p = 0.166, d = 0.97) (Fig. 2A). *PGC-1α1* mRNA increased from Pre to +3h in ISC (p = 0.003, d = 1.36), but was not significantly altered in CON (p = 0.649, d = 0.29) or in HYP (p = 0.702, d = 0.32). The increase in ISC was greater compared to that in CON (p = 0.017, d = 1.45) and in HYP (p = 0.040, d = 1.21) (Fig. 2B). *PGC-1α4* mRNA increased from Pre to +3h in ISC (p = 0.002, d = 1.25), but was not significantly increased in CON (p =0.333, d = 1.12) or in HYP (p = 0.272, d = 0.86). The increase in ISC was greater than in CON (p = 0.037, d = 1.15) and in HYP (p = 0.057, d= 0.77) (Fig. 2C).

**Figure 2.**
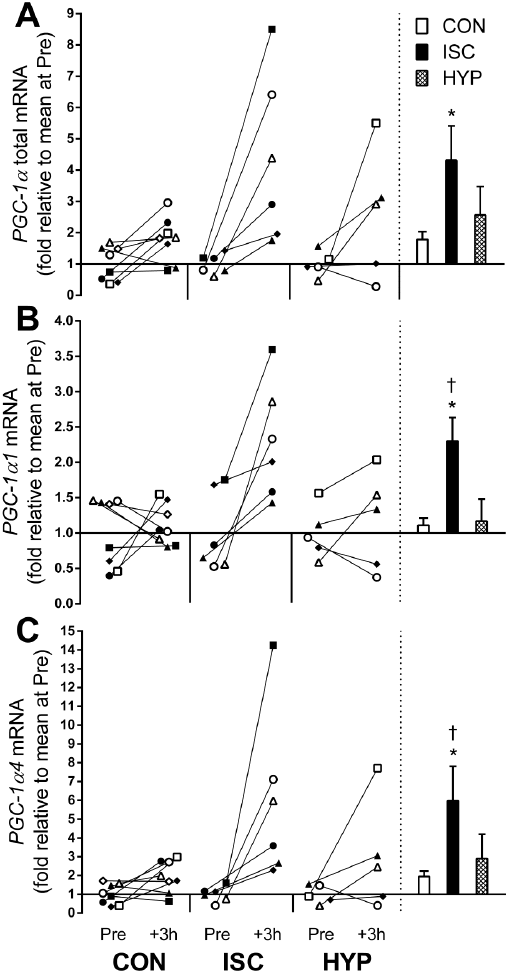
PGC-1α total and -isoform mRNA responses to repeated exercise with ischaemia or in systemic hypoxia. A) *PGC-1α* total, B) *PGC-1α_1_*, and C) *PGC-1α_4_*, mRNA content. Individual changes from before (Pre) to 3 h after exercise (+3h) are displayed on the left with each symbol representing one participant across trials and figures. On the right are bars representing mean (± SEM) changes relative to Pre in the dominant leg for exercise alone (CON, white), with ischaemia (ISC occluded, black), or in systemic hypoxia (HYP, meshed). ^*^ p ≤ 0.05, different from Pre; † p ≤ 0.05, different from CON and HYP.

### Na^+^,K^+^-ATPase (NKA) and FXYD1 gene transcripts

*NKAα1* mRNA remained unchanged in ISC (p = 0.799, d = 0.44), CON (p = 0.648, d = 0.54), and in HYP (p = 0.557, d = 0.32) (Fig. 3A). *NKAα _2_* mRNA increased from Pre to +3h in ISC (p= 0.050, d = 0.90) and in HYP (p = 0.004, d = 0.61), but there was no significant change in CON (p = 0.089, d = 1.07) (Fig. 3B). *NKAα_3_* mRNA was not significantly altered in all conditions (p ≥ 0.169, d = 0.57, 0.63 and 0.58 in ISC, CON and HYP, respectively) (Fig. 3C). *NKAβ_1_* mRNA increased in HYP (p = 0.027, d = 0.68), but there was no significant change in ISC (p= 0.268, d = 0.79) or in CON (p = 0.539, d = 1.19) (Fig. 4A). *NKAβ_2_* mRNA was not significantly altered in all conditions (p ≥ 0.276, d = 0.24, 0.40 and 0.24 in ISC, CON and HYP, respectively) (Fig. 4B). *NKAβ_3_* mRNA was also not significantly changed all conditions (ISC: p = 0.909, d = 0.34; CON: p = 0.859, d = 0.55; HYP: p = 0.328, d = 0.52) (Fig. 4C). FXYD1 mRNA increased in ISC (p = 0.016, d = 1.10). This increase was greater than in CON (p = 0.039, d = 1.05). There were no changes in the remaining conditions for this gene (CON: p = 0.915, d = 0.20; HYP: p = 0.305, d = 0.32) (Fig. 4D).

**Figure 3.**
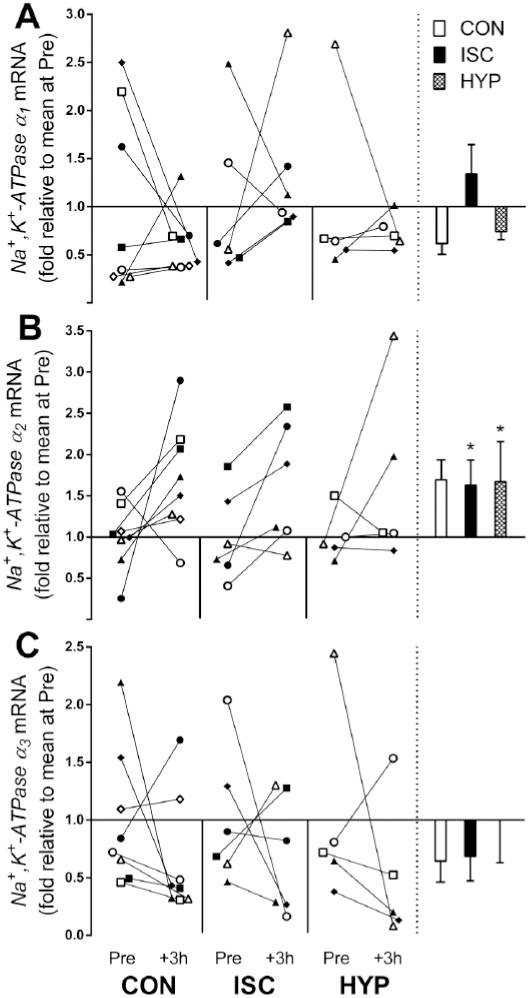
NKA-α-isoform mRNA responses to repeated exercise with ischaemia or in systemic hypoxia. A) *α*_*1*_, B) *α*_*2*_, and C) *α*_*3*_, mRNA content. Individual changes from before (Pre) to 3 h after exercise (+3h) are displayed on the left with each symbol representing one participant across trials and figures. On the right are bars representing mean (± SEM) changes relative to Pre in the dominant leg for exercise alone (CON, white), with ischaemia (ISC occluded, black), or in systemic hypoxia (HYP, meshed). ^*^ p ≤ 0.05, different from Pre.

**Figure 4.**
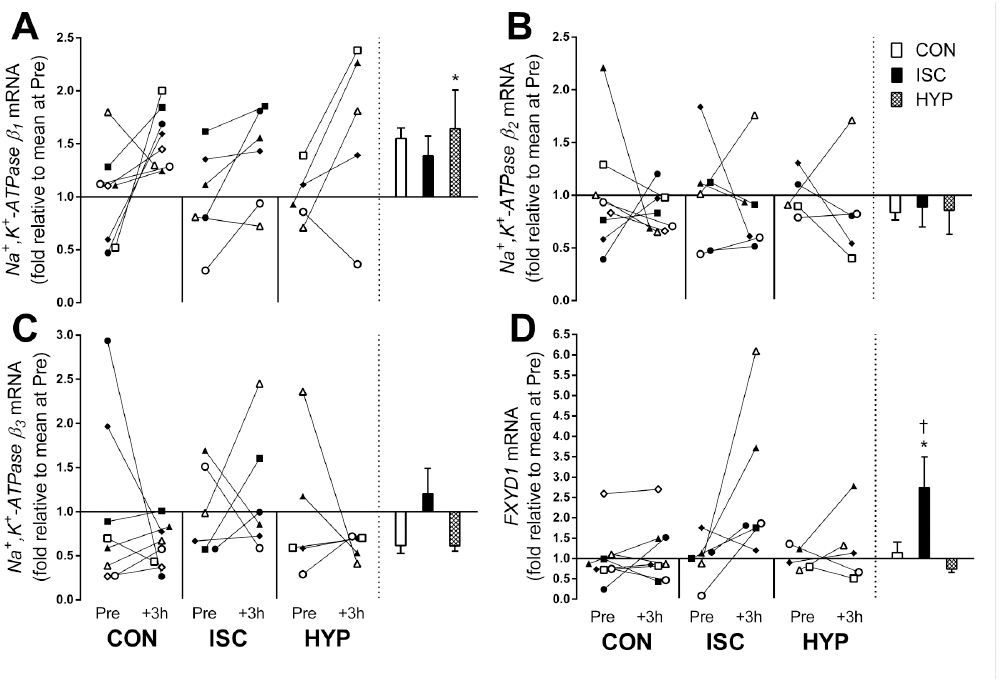
NKA-β-isoform and FXYD1 mRNA responses to repeated exercise with ischaemia or in systemic hypoxia. A) *β*_*1*_, B) *β*_*2*_, C) *β*_*3*_, and D) FXYD1, mRNA content. Individual changes from before (Pre) to 3 h after exercise (+3h) are displayed on the left with each symbol representing one participant across trials and figures. On the right are bars representing mean (± SEM) changes relative to Pre in the dominant leg for exercise alone (CON, white), with ischaemia (ISC occluded, black), or in systemic hypoxia (HYP, meshed). ^*^ p < 0.05, different from Pre; † p < 0.05, different from CON.

### Muscle deoxygenation and oxidative stress markers

Muscle deoxygenation, as assessed by deoxygenated haemoglobin concentration (muscle HHb), was higher (p < 0.05) during exercise in ISC and HYP, relative to CON, except for the seventh bout of exercise (p > 0.05). During the recovery from the first, fourth and fifth bout, muscle HHb was higher in ISC, relative to CON (p < 0.05). During the recovery from the second bout, muscle HHb was higher in ISC and HYP, compared to CON (p > 0.05; Fig. 5A). No significant differences were detected between ISC and HYP at any time point (p > 0.05). *Catalase* mRNA content increased significantly in ISC (p = 0.024, d = 0.70), but there was no significant changes in CON (p = 0.881, d = 0.34) or in HYP (p = 0.505, d = 0.15). *HSP70* mRNA increased in ISC (p = 0.057, d = 0.86), but there was no significant changes in CON (p= 0.669, d = 0.18) or in HYP (p = 0.176, d = 0.95).

**Figure 5.**
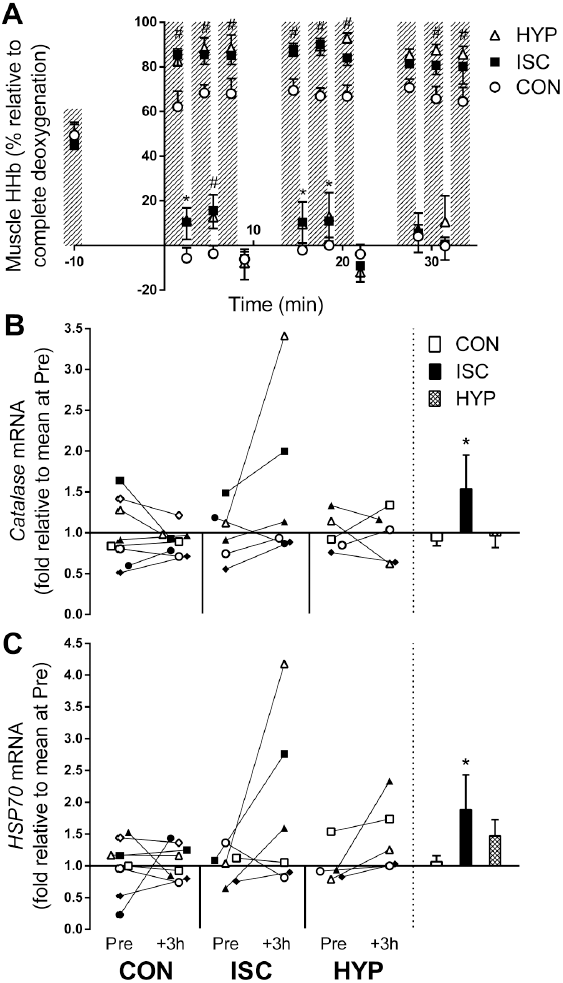
Changes in muscle deoxygenation and oxidative stress markers in response to repeated exercise with ischaemia or in systemic hypoxia. A) Muscle deoxygenaton (i.e. deoxygenated haemoglobin, Muscle HHb) as assessed by near-infrared spectroscopy during exercise alone (CON, o), with ischaemia (ISC, ■), or in systemic hypoxia (HYP, Δ). Hashed bars represent exercise bouts. # p ≤ 0.05, ISC and HYP different from CON. ^*^ p ≤ 0.05, ISC different from CON. B) Catalase, and C) heat-shock protein 70 (HSP70), mRNA expression. Individual changes from before (Pre) to 3 h after exercise (+3h) are displayed on the left with each symbol representing one participant across trials and figures. On the right are bars representing mean (± SEM) changes relative to Pre in the dominant leg for exercise alone (CON, white), with ischaemia (ISC occluded, black), or in systemic hypoxia (HYP, meshed). Data are means ± SEM. ^*^ p ≤ 0.05, different from Pre.

### Muscle metabolites

Muscle ATP remained unchanged in all conditions (p > 0.05; Fig. 6A). Muscle lactate increased in ISC and in HYP, but not in CON (p > 0.05). The increase in ISC and HYP was greater than CON (p < 0.05; Fig. 6B).

**Figure 6.**
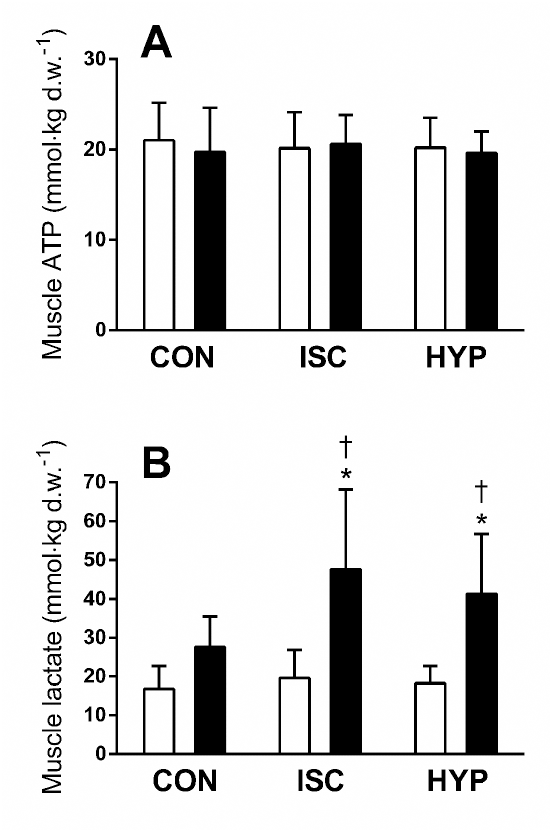
Changes in muscle ATP and lactate concentration in response to repeated exercise with ischaemia (ISC) or in systemic hypoxia (HYP). A) ATP, and B) lactate concentration before (Pre, white) and immediately after exercise (+0h, black). Data are means ± SEM. ^*^ p < 0.05, different from Pre. † p < 0.05, different from CON.

### Venous blood lactate, pH and potassium ion concentration

In ISC, blood lactate concentration ([lac^-^]) increased (p < 0.05) after the third bout of exercise and remained elevated throughout the trial compared to rest. Blood [lac-] was higher (p < 0.05) in ISC than in CON after the third bout, the fifth to ninth bout, and after 3 min of recovery. In HYP, blood [lac^-^] increased (p < 0.05) after the third, fifth, sixth, eighth and ninth bout and in recovery, compared to rest. Blood [lac^-^] was higher (p < 0.05) in HYP than in CON after the third, fifth and ninth bout of exercise. In CON, blood [lac^-^] remained unchanged throughout the trial, compared to rest (p > 0.05).

In ISC, blood pH dropped (p < 0.05) following the first bout of exercise and remained lower (p< 0.05) compared to rest throughout the trial, and 3 min into recovery, but returned to resting level after 6 min of recovery (p > 0.05). The drop in pH in ISC was lower, relative to CON, after the sixth, before the seventh, and after the eighth and ninth, bout relative to CON (p < 0.05). In HYP, blood pH was lower, compared to rest, following the third, fifth, and sixth bout, and before the seventh bout, but returned to resting level after the seventh bout, from where it remained unchanged. The drop in pH in HYP was lower following the sixth and before the seventh bout, relative to CON (p < 0.05). In CON, blood pH remained unchanged throughout the trial (p > 0.05).

In all trials, blood potassium ion concentration ([K^+^]) increased after warm-up, and after the first to eighth bout of exercise, compared to rest. In CON, blood [K+] was also elevated (p < 0.05) after the ninth bout of exercise, relative to rest. Compared to CON, blood [K^+^] was lower (p < 0.05) in ISC 4 min into recovery from the third bout, and 6 min into recovery from the ninth bout of exercise, with no differences at other time points (p > 0.05), nor between HYP and CON at all time points (p > 0.05).

### AMPK and ACC protein abundance and phosphorylation

Representative blots for AMPK and ACC are shown in Fig. 8A.

**Figure 7.**
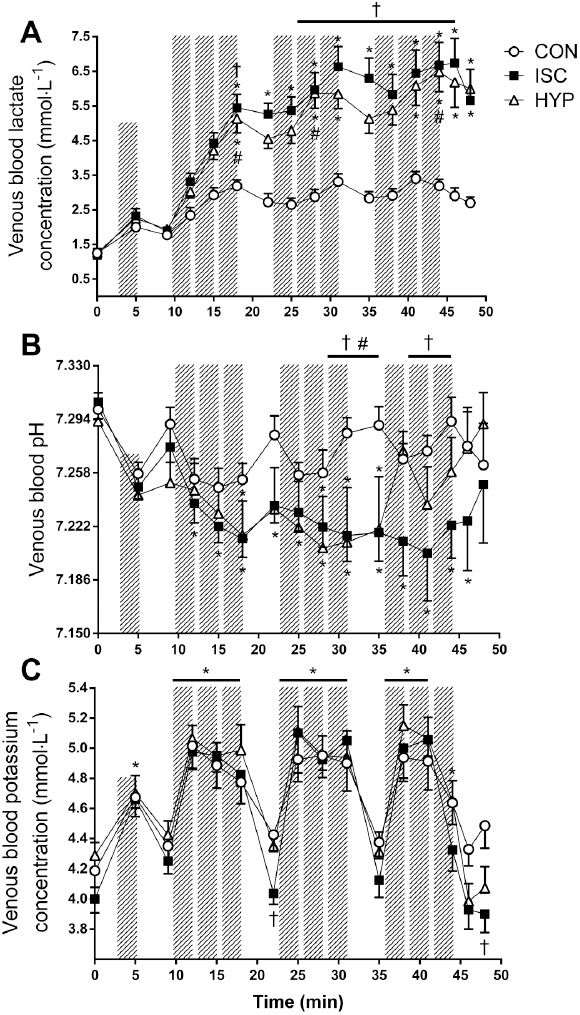
Changes in venous blood lactate, pH, and potassium ion (K^+^) concentration in response to exercise with ischaemia or in systemic hypoxia. A) Lactate, B) pH, and C) K^+^ concentration during exercise alone (CON, o), with ischaemia (ISC, ■), or in systemic hypoxia (HYP, Δ). Hashed bars represent exercise bouts. Data are means ± SEM. ^*^ p < 0.05, different from rest; † p < 0.05, ISC different from CON; # p < 0.05, HYP different from CON.

**Figure 8.**
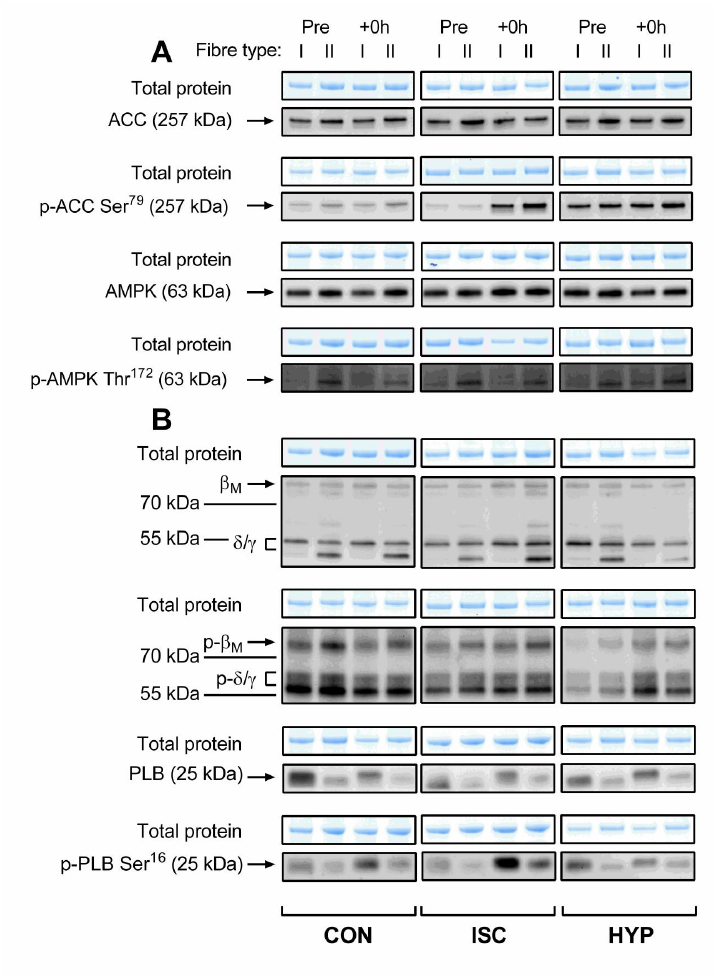
Representative blots for AMPKα, ACC, CaMKII, and phospholamban (PLB) protein abundance and phosphorylation in type I and II human skeletal muscle fibres. A) AMPK and ACC, and B) CaMKII and PLB, protein abundance and phosphorylation in the dominant leg in response to exercise alone (CON), with ischaemia (ISC occluded), or in systemic hypoxia (HYP) before (Pre) and immediately after (+0h) exercise. Total protein was determined in each lane from the stain-free gel images obtained after electrophoresis. CaMKII isoforms (βM and σ/γ) are indicated in B.

In HYP, α-AMPK protein abundance decreased (p = 0.023) from Pre to +0h in type I, but did not change significantly in type II (p = 0.112) fibres. In ISC and CON, α-AMPK protein abundance was not significantly altered in both fibre types (p ≥ 0.423). The abundance was significantly higher in both fibre types in ISC relative to HYP at +0h (p ≤ 0.002) (Fig. 9A). The α-AMPK protein abundance was significantly higher in type II vs. type I fibres in all conditions (main effect of fibre type, p ≤ 0.027). In HYP, the phosphorylation of α-AMPK at Thr172 relative to total α-AMPK protein abundance (α-AMPK Thr^172^/α-AMPK) increased significantly in type I (p = 0.003), but not in type II (p = 0.558) fibres. In ISC and CON, there was no significant change in α-AMPK Thr^172^/α-AMPK in both fibre types (p ≥ 0.112). The α-AMPK Thr^172^/α-AMPK was significantly higher in type II vs. type I fibres in all conditions (main effect of fibre type, p≤ 0.017) (Fig. 9B).

**Figure 9.**
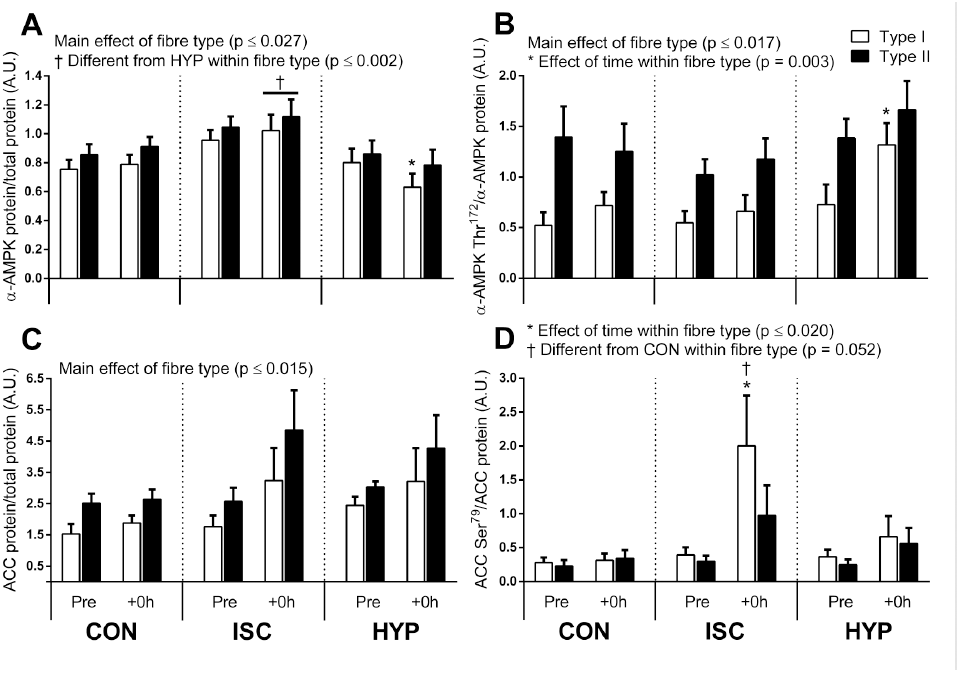
Changes in AMPKα and ACC protein abundance and phosphorylation in type I and II human skeletal muscle fibres in response to repeated exercise with ischaemia or in systemic hypoxia. A) AMPKα protein, B) AMPKα phosphorylation at Thr^172^ normalised to AMPKα protein, C) ACC protein, and D) ACC phosphorylation at Ser^79+^ normalised to ACC protein in type I (white bars) and type II (black bars) fibres before (Pre) and immediately after (+0h) exercise. Data are means ± SEM. ^*^ p < 0.05, different from rest within condition and fibre type;† p ≤ 0.05, ISC different from HYP in A, and from CON in D.

ACC protein abundance was not significantly altered in both fibre types in all conditions (p ≥ 0.168), and overall it was higher in type II vs. type I fibres (main effect of fibre type, p ≤ 0.015; Fig. 9C). The phosphorylation of ACC at Ser^79^ to total ACC protein (ACC Ser^79^/ACC) increased significantly from Pre to +0h in type I fibres in ISC (p ≤ 0.020), with the increase being significantly higher relative to CON (p = 0.052). In the same condition, there was no significant change in ACC Ser^79^/ACC in type II fibres (p = 0.260). No changes in ACC Ser^79^/ACC occurred in CON and HYP (p ≥ 0.213) (Fig. 9D).

### CaMKII and phospholamban (PLB) protein abundance and phosphorylation

Representative blots for CaMKII and PLB are shown in Fig. 8B.

CaMKII protein abundance did not change significantly in both fibre types in all conditions (p≥ 0.108). In ISC and CON, it was significantly higher in type I vs. type II fibres (main effect of fibre type, p ≤ 0.038), but not significantly different between fibre types in HYP (p = 0.200) (Fig. 10A). Phosphorylation of CaMKII at Thr^287^ to total CaMKII protein (CaMKII Thr^287^/CaMKII) decreased significantly in type II fibres in CON (p = 0.023), and tended to decrease in ISC (p= 0.056), but did not change significantly in HYP (p = 0.746). No significant changes in CaMKII Thr^287^/CaMKII occurred in type I fibres in all conditions (p ≥ 0.213). CaMKII Thr^287^/CaMKII was significantly higher in type II vs. type I fibres in all conditions (main effect of fibre type, p ≤ 0.023) (Fig. 10B).

**Figure 10.**
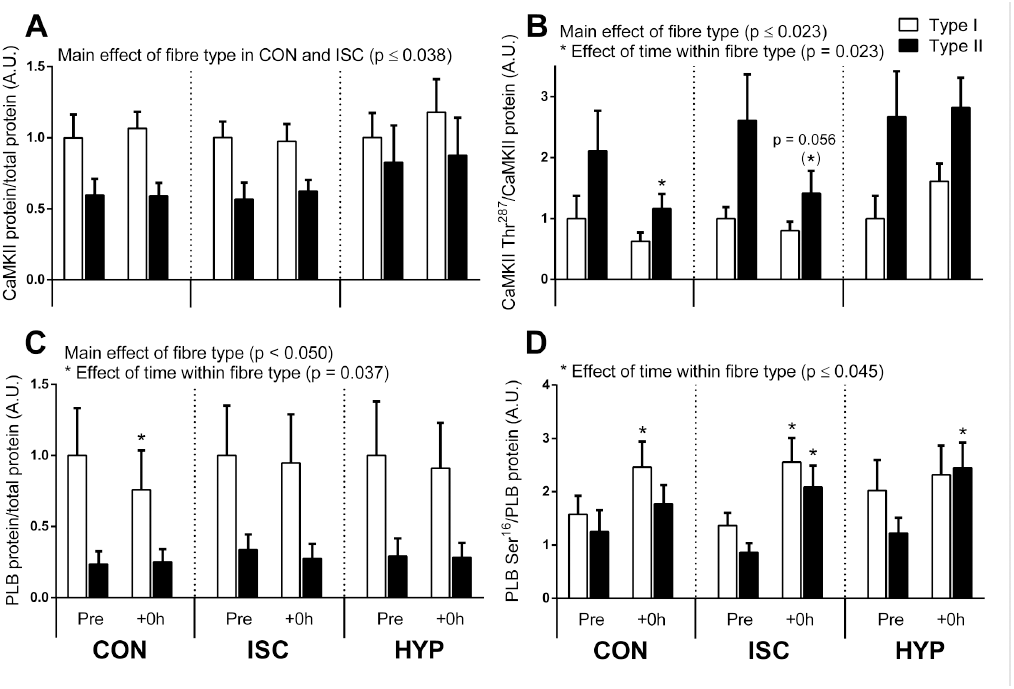
Changes in CaMKII, and phospholamban (PLB) protein abundance and phosphorylation in type I and II human skeletal muscle fibres in response to repeated exercise with ischaemia or in systemic hypoxia. A) CaMKII protein, B) CaMKII phosphorylation at Thr^287^ normalised to CaMKII protein, C) PLB protein, and D) PLB phosphorylation at Ser^16^ normalised to PLB protein in type I (white bars) and type II (black bars) fibres before (Pre) and immediately after (+0h) exercise. Data are means ± SEM. ^*^p < 0.05, different from rest within condition and fibre type.

In type I fibres, PLB protein abundance decreased significantly in CON (p = 0.037), whereas it did not change significantly in ISC (p = 0.786) or in HYP (p = 0.428) in the same fibre type. The PLB abundance did not change significantly in type II fibres in all conditions (p ≥ 0.288). The PLB abundance was lower in type II vs. type I fibres (main effect of fibre type, p ≤ 0.050) (Fig. 8C). The phosphorylation of PLB at Ser^16^ to total PLB protein (PLB Ser^16^/PLB) increased significantly in type I fibres in CON (p = 0.023) and in ISC (p = 0.010), but it remained unchanged in HYP in the same fibre type (p = 0.412). In type II fibres, PLB Ser^16^/PLB increased in ISC (p ≤ 0.026) and in HYP (p = 0.025), but did not change significantly in CON (p = 0.350) (Fig. 8D).

## Discussion

The main novel findings, which are summarised in Fig. 11, were that repeated bouts of ischaemic exercise substantially elevated the mRNA content of PGC-1α (~4.3 fold), PGC-1α1 (~2.3 fold) and PGC-1α4 (~6 fold), and of the NKA regulatory subunit gene, FXYD1 (~2.7 fold), in trained human skeletal muscle. These changes coincided with increased levels of oxidative stress markers (*catalase* and *HSP70* mRNA, 1.5-1.9 fold), and were temporally preceded by elevated (~1-2 fold) ACC phosphorylation (Ser^79^) in both muscle fibre types. These results suggest the effect of ischaemia on exercise-induced changes in *PGC-1α* and *FXYD1* mRNA levels was mediated, at least in part, through potentiation of ROS formation and activation of associated signalling networks, of which AMPK could be one (contributing) pathway (Irrcher *et al.*, 2009). Despite a similar exercise-induced reduction in muscle oxygen content in the ischaemic and hypoxic trials (ISC vs. HYP, Fig. 5), we found no impact of exercising in systemic hypoxia on either the mRNA content of PGC-1α isoforms and FXYD1, or on markers of oxidative stress and ACC phosphorylation. These findings support that perturbations in redox homeostasis, rather than a critical threshold of muscle hypoxia *per se*, were key to the transcriptional induction of PGC-1α isoforms and FXYD1 in response to the repeated-ischaemic exercise (Stoner *et al.*, 2007).

**Figure 11.**
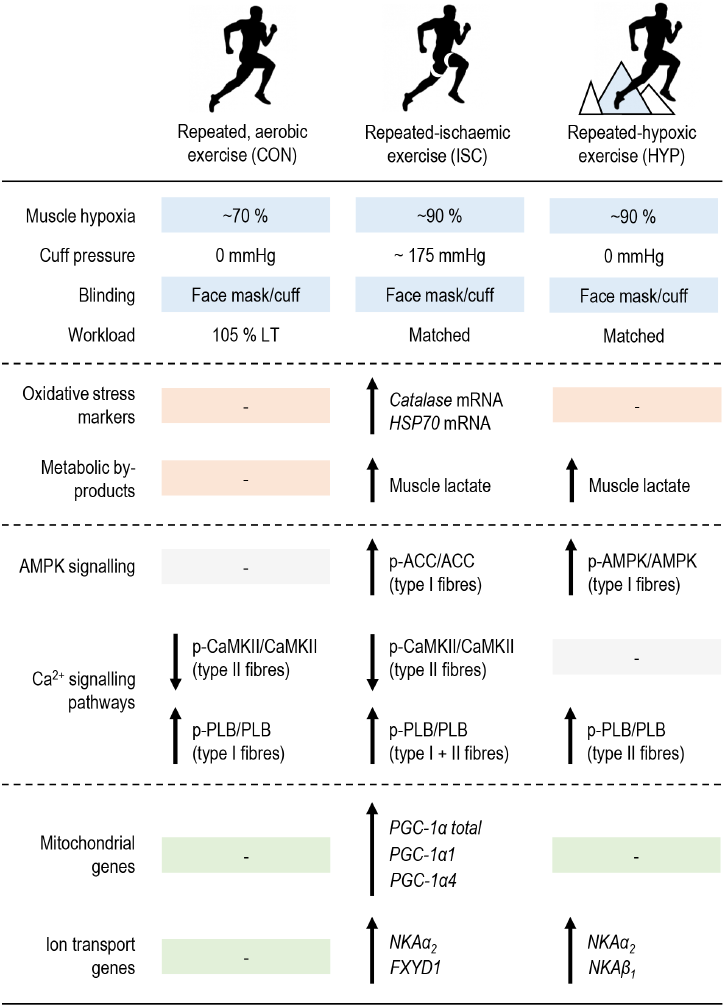
Summary of key findings. The effect of aerobic interval (CON), repeated-ischaemic (ISC) and –hypoxic exercise (ISC) on the mRNA content of PGC-1α (total, *1α1, 1α4*) and Na^+^,K^+^-ATPase (*NKAα_1-3_*, *NKAβ_1-3_*, *FXYD1*) isoforms, oxidative stress markers (*catalase* and *heat-shock protein 70, HSP70*, mRNA content), muscle hypoxia (i.e. deoxygenated haemoglobin as measured by near-infrared spectroscopy), lactate concentration, and AMPK and CaMKII signalling in the skeletal muscle of trained men. “p-“ denotes phosphorylation; ACC, Acetyl-CoA carboxylase; AMPK, 5' AMP-activated protein kinase; CaMKII, Ca2+/calmodulin-dependent protein kinase II; PLB, phospholamban; LT, lactate threshold.

### Repeated-ischaemic exercise increases PGC-1α total and -isoform mRNA levels in the skeletal muscle of trained men

In a previous human study, a decrease of ~15-20 % in muscle blood perfusion (Eiken & Bjurstedt, 1987) augmented the rise (~5.5 fold) in muscle *PGC-1α* (total) mRNA content in response to knee-extensor exercise (45 min at 26 % of one-leg peak load) (Norrbom *et al.*, 2004). In accordance, in the present study, ischaemic exercise induced a 4.3-fold rise (i.e. 2.5-fold greater increase vs. CON) in total *PGC-1α* expression (Fig. 2). Furthermore, the mRNA levels of both the *PGC-1α1* (2.5-fold) and *PGC-1α4* (6.0 fold) were elevated only after the ischaemic exercise. Our results highlight for the first time the effectiveness of this training modality to enhance the exercise-induced response of PGC-1α-isoform transcripts in humans.

Oral consumption of antioxidants (i.e. vitamin C and E) have been shown to blunt the exercise-induced increase in *PGC-1α* mRNA content in the skeletal muscle of young men (Ristow *et al.*, 2009), suggesting ROS could be important for the upregulation of *PGC-1α* mRNA content in human muscle. Accordingly, the increases in PGC-1α-isoform mRNA levels in ISC coincided with elevated *catalase* and *HSP70* mRNA levels, both shown to be specifically induced by hydrogen peroxide (H_2_O_2_) in myocytes (Kemp *et al.*, 2003; Zhong *et al.*, 2011). These data provide the first evidence in humans that facilitated ROS production may be one mechanism underlying the increases in *PGC-1α1* and *-1α4* mRNA transcript levels. In support, several *in vitro* studies have suggested ROS may be implicated in *PGC-1α* transcription. For example, in C_2_C_12_ cells, incubation with H_2_O_2_ elevated both *PGC-1α* mRNA content (1.4 fold) and promoter activity (Irrcher *et al.*, 2009), whereas pre-incubation with the ROS scavenger, N-acetylcysteine (NAC) abolished the H_2_O_2_-evoked rise in promoter activity (Zhang *et al.*, 2014) and mRNA content (Irrcher *et al.*, 2009). Together, our results suggest that the effect of ischaemia was mediated, in part, by promotion of ROS generation in skeletal muscle.

### Association between increases in PGC-1α mRNA transcripts and ACC phosphorylation

In line with our second working hypothesis, the ischaemia-evoked increases in *PGC-1α* mRNA transcripts were preceded by elevated ACC phosphorylation, indicative of higher AMPK activity (Chen *et al.*, 2000). This suggests AMPK could be one signalling kinase through which the initial cellular signals are transduced to regulate the content of multiple *PGC-1α* mRNA transcripts in human muscle. This hypothesis is consistent with data from cell culture studies. In one of these studies, treatment of C2C12 cells with the AMPK activator, 5-aminoimidazole-4carboxamide-1-b-D-ribofuranoside (AICAR), raised *PGC-1α* mRNA content (2.2 fold) coincidently with augmented *PGC-1α* promoter activity (3.5 fold). In that study, several AICAR-sensitive *PGC-1α* promoter sites were identified (Irrcher *et al.*, 2008), suggesting AMPK may directly facilitate *PGC-1α* transcription. In a more recent experiment, transcriptional activation of *PGC-1α* by ROS coincided with promoted AMPK activation (Irrcher *et al.*, 2009). Thus, *in vitro*, ROS-induced activation of *PGC-1α* transcription could be mediated in part via AMPK.

The coincident increases in oxidative stress markers, ACC phosphorylation, and PGC-1α- isoform mRNA content with ischaemia in the present study underline that a similar signal transduction axis involving ROS and AMPK may exist in humans. However, factors other than ROS may, in part, account for the higher ACC phosphorylation with ischaemia. For example, circulating norepinephrine can stimulate AMPK activity in skeletal muscle cells via α_1_-adrenoceptors (Hutchinson & Bengtsson, 2006), and ischaemia has been shown to exacerbate exercise-induced increases in circulating norepinephrine concentration (Sundberg, 1994). Moreover, the increase in ACC phosphorylation was most prominent in type I fibres (Fig. 9D), consistent with an accelerated glycogenolysis in this fibre type during repeated contractions with circulatory occlusion (250 mmHg) (Greenhaff *et al.*, 1993). These findings support that the mRNA responses to ischaemic exercise could be fibre type-dependent. Future work is required to examine this further by measuring PGC-1α-isoform mRNA and AMPK activity in different muscle fibre types. Inconsistent with the changes in ACC phosphorylation, phosphorylation of AMPKα remained unaltered in ISC and CON, and as such, was dissociated from the changes in *PGC-1α* transcript levels. The latter finding is in line with previous observations in humans (Brandt *et al.*, 2016; Taylor *et al.*, 2016). This dissociation of AMPK and ACC phosphorylation is likely explained by the extent to which phosphorylation of different AMPK heterotrimer complexes affects muscle AMPK activity (Birk & Wojtaszewski, 2006).

### Increases in PGC-1α-isoform mRNA content are unrelated to modulation of CaMKII phosphorylation in human muscle fibre types

In myocytes *in vitro*, inhibition of CaMK has been reported to abolish Ca^2+^induced increases in PGC-1α expression *(Ojuka et al., 2003)*. Thus, we explored the association between changes in *PGC-1α* mRNA levels and those of CaMKII phosphorylation at Thr^287^, which in human muscle has been correlated with CaMKII autonomous activity (r^2^= 0.884) (Rose *et al.*, 2006). A novel observation was that CaMKII phosphorylation decreased in type II fibres only in ISC (p = 0.056) and CON. This fibre type-specific modulation of CaMKII phosphorylation was abolished by systemic hypoxia, indicating that decreased arterial oxygen saturation may affect contraction-stimulated Ca^2+^signalling in type II human muscle fibres. This effect of hypoxia, however, was dissociated from alterations in *PGC-1α* levels. Changes in CaMKII phosphorylation, regardless of fibre type, were unrelated to those of *PGC-1α* in the ischaemic-exercised leg. Together, these data imply that promotion of human muscle PGC-1α-isoform mRNA levels does not require changes in CaMKII autonomous activity *per se*, although our observations are limited to immediately post exercise. In mouse fast-twitch skeletal muscle fibres, CaMKII partly controls SR Ca^2+^ release, but not SR Ca^2+^ uptake (Tavi *et al.*, 2003). It may therefore be speculated whether the decrease in CaMKII phosphorylation in type II fibres in ISC and CON (Fig. 10) may have reflected a reduced SR Ca^2+^ leak, although this cannot be resolved from the present data alone. Nevertheless, ischaemia may have affected type-II fibre SR Ca^2+^ uptake, as ischaemia invoked an increase in PLB phosphorylation in this fibre type (Fig. 10D). In agreement, relieving the inhibition of PLB by phosphorylation at Ser16 promotes SR Ca^2+^ uptake in both fibre types, with the effect being most pronounced in predominantly glycolytic fibres (Briggs *et al.*, 1992). A similar fibre type-dependent effect on PLB phosphorylation was evident with systemic hypoxia. Based on our observations, modulation of SR Ca^2+^ kinetics may not have been a primary mechanism underlying the effect of ischaemia on *PGC-1α* mRNA levels in the present study.

### Increases in human muscle PGC-1α-isoform mRNA content are unrelated to muscle hypoxia

Unlike ischaemia, exercise alone was without impact on the levels of the investigated *PGC-1α* transcripts. This may be explained by a low relative exercise intensity (105 % LT) given the high training status of our participants, the moderate running speed (~12 km·h-^1+^), and the positive relation between exercise intensity and exercise-induced increases in muscle *PGC-1α* mRNA content previously reported (Egan *et al.*, 2010; Nordsborg *et al.*, 2010b). However,absence of an effect of exercise alone was unlikely related to muscle metabolic stress *per se*. Accordingly, we found no effect of systemic hypoxia on *PGC-1α* levels, despite exacerbated metabolic by-product accumulation comparable to that induced by ischaemic exercise (Fig. 6 and 7). This is consistent with a previous human study, in which contraction-stimulated increases in leg muscle *PGC-1α* mRNA content was unaffected by promoted systemic levels of epinephrine and lactate invoked by simultaneous arm exercise (Brandt *et al.*, 2016). The present observation of unaltered *PGC-1α* levels after exercise in systemic hypoxia agrees with a previous human study that reported unchanged *PGC-1α* mRNA content after a single exercise session at simulated altitude (3000 m) in recreationally active men (Slivka *et al.*, 2014). Since deoxygenated HHb was similar during exercise with ischaemia and in systemic hypoxia (Fig. 5A), this lack of a hypoxic effect may not be ascribed to cellular hypoxia *per se*. As such, the severity of exercise-induced muscle hypoxia was not decisive for the ischaemia-mediated increases in *PGC-1α* transcript levels in the present study. Given the comparable fluctuations in muscle lactate and blood metabolites in HYP and ISC (Fig. 6 and 7), inadequate metabolic by-product accumulation may also not account for the lack of a hypoxic effect. As we detected no change in oxidative stress markers with hypoxia, it may rather be explained by insufficient ROS accumulation, although a single factor may not fully account for this absence of effect given the cross-talk and redundancy of ionic and metabolic perturbations, and ROS formation (Kourie, 1998).

### Possible involvement of ROS in the regulation of FXYD1 mRNA content in human muscle

Another novel result was that *FXYD1* mRNA content increased (2.7 fold) due to ischaemia (Fig. 4D). Despite similar increases in deoxygenated HHb between HYP and ISC (Fig. 5A), systemic hypoxia was without effect on *FXYD1* expression. Thus, cellular hypoxia was also unlikely a primary stimulus for the ischaemia-induced increase in *FXYD1*. Nor may the increase be related to the severity of metabolic stress, as changes in muscle lactate and phosphocreatine were similar in ISC and HYP (Fig. 6). The rise in *FXYD1* mRNA expression, however, was accompanied by increases in markers of oxidative stress (Fig. 5). Therefore, accumulation of ROS may be an important mechanism underlying the effect of ischaemia on *FXYD1* transcription. Accordingly, FXYD1 overexpression protected myocytes against ROS-induced NKA dysfunction (Liu *et al.*, 2013), suggesting a ROS-protective effect of elevated FXYD1 content. It has also been established in cell culture that AMPK can be activated by ROS (Irrcher *et al.*, 2009), and this regulates *FXYD* transcription in mouse glycolytic skeletal muscles (Nilsson *et al.*, 2006). In accordance, the increases in *FXYD1* and oxidative stress markers were paralleled by elevated ACC phosphorylation (Fig. 9). Therefore, regulation of human muscle *FXYD1* expression could involve both ROS production and AMPK activation. Measurement of AMPK activity, along with *FXYD1* mRNA content, should be performed in future studies to verify these initial results. In addition, alterations in *FXYD1* expression were dissociated from those of CaMKII and PLB phosphorylation (Fig. 10), suggesting transcriptional induction of *FXYD1* mRNA content in human muscle does not require alterations in CaMKII autonomous activity.

### Increases in human muscle NKA α_1_ and β_3_ mRNA content are unrelated to oxidative stress and metabolic perturbations

In the present study, *NKAα_1_* and -*β*_*3*_ mRNA content was unaffected by exercise with ischaemia, in systemic hypoxia, or by exercise alone (Fig. 3A and 4C), despite substantial differences amongst conditions in changes in markers of oxidative stress, muscle lactate, and blood metabolites. As such, the level of these mRNA transcripts are neither influenced by the nature of metabolic and ionic fluctuations, nor by the degree of oxidative stress, in human skeletal muscle. In support, raising the metabolic stress by performing simultaneous arm exercise was without effect on the increases in *α*_*1*_ and *β*_*3*_ mRNA content after isolated knee-extensions (Nordsborg *et al.*, 2005). Based on the individual changes for these isoforms in the present study (Fig. 3A and 4C), they seem to be regulated similarly independent of condition. For example, the same two individuals that decreased their *α*_*1*_ mRNA content in ISC also reduced their *β*_*3*_ expression in the same condition. Accordingly, we have recently shown parallel increases in only *α*_*1*_ and *β*_*3*_ mRNA levels following sprint interval exercise (Christiansen et al. 2017, *unpublished*). In another human study, *NKAα_1_* and *-β_3_* were the only transcripts of those investigated (*α*_*1-3*_ and *β*_*1-3*_) that remained unaltered in response to 45 min cycling at 71 % VO_2ma_x (Murphy *et al.*, 2008). Together, these results highlight that α_1_ and β_3_ are likely regulated at the mRNA level by the same cellular signals, different from those important to changes in the mRNA expression of other NKA isoforms (e.g. compare the individual changes for *α*_*1*_ and *α*_*2*_; Fig. 3A and 3B).

### Regulation of other NKA-isoform mRNA transcripts by repeated-ischaemic exercise

The NKA α_2_ isoform is limiting for a muscle’s contractile performance (Radzyukevich *et al.*, 2013) and forms up to 90 % of NKA complexes in adult rat skeletal muscles (Orlowski & Lingrel, 1988). Together with the β_1_ isoform, it forms the largest NKA pool in this tissue. Understanding the cellular signals that regulate α_2_ and β_1_ expression is therefore fundamental. In the present study, significant increases in the mRNA content of *NKAα_2_* and *-β_1_* were evident in HYP and ISC, and in HYP alone, respectively, whereas CON was without effect on these genes. This suggests a hypoxic and/or a metabolically perturbed cellular milieu could be beneficial for increasing the mRNA content of these isoforms in human muscle. However, taking into account the individual values for *α*_*2*_ and *β*_*1*_ (Fig. 3B), there appears no obvious difference in the changes in both *α*_*2*_ and *β*_*1*_ levels between CON, ISC, and HYP. This raises the possibility of a statistical type II error (in CON for *α2* and in CON and ISC for *β*_*1*_). This precludes us from unequivocally interpreting the present data related to these isoforms. More research appears required to elaborate on the idea that cellular hypoxia and/or metabolic stress *per se* are important stimuli for increases in α_2_-and β_1_-isoform expression with contractile activity. In addition, no alterations in the levels of *NKAα_3_* and -*β*_*2*_ transcripts were found for any condition (Fig. 3 and 4). We have previously observed no change in *α*_*3*_ mRNA content after repeated-intense exercise of short duration (Christiansen et al. 2017, in review, FASEB). Conversely, exercise-induced increases in *α*_*3*_ expression have been reported in other human studies. In these studies, the changes in *α*_*3*_ mRNA occurred at exercise termination, with the level returning to basal state after 3 h of recovery (Murphy *et al.*, 2004; Aughey *et al.*, 2007; Murphy *et al.*, 2008). Thus, our time point of mRNA measurement may have been decisive for the present outcome, and as such is a limitation of the present study. The effect of exercise on *β*_*2*_ mRNA expression is controversial with human studies reporting both increased, decreased, or unchanged, expression 3 h into recovery. In the present study, the level of *β*_*2*_ mRNA remained unchanged. The reason for these conflicting findings is not clear (Christiansen et al. 2017, *unpublished*), and further mechanistic studies are needed to understand how β_2_ expression is regulated in human muscle.

### Conclusion and perspectives

In summary, repeated bouts of ischaemic exercise potently raised the mRNA levels of PGC-1α, −1α1 and −1α4, and of the NKA regulatory subunit, FXYD1, in skeletal muscle of trained men. The molecular mechanisms likely involved were promoted ROS generation and AMPK activation. The effect of ischaemia was, however, unrelated to the severity of muscle hypoxia, lactate accumulation, and fibre type-specific modulation of CaMKII signalling. Thus, repeated-ischaemic exercise is a potent strategy to enhance the muscle’s gene response associated with mitochondrial biogenesis and K^+^ handling in trained individuals. Based on this work, future studies should evaluate whether the muscle’s capacity for oxidative ATP generation and potassium ion regulation would be improved by a period of repeated-ischaemic training.

## References

Abe T, Fujita S, Nakajima T, Sakamaki M, Ozaki H, Ogasawara R, Sugaya M, Kudo M, Kurano M, Yasuda T, Sato Y, Ohshima H, Mukai C & Ishii N. (2010). Effects of Low-Intensity Cycle Training with Restricted Leg Blood Flow on Thigh Muscle Volume and VO2MAX in Young Men. Journal of sports science & medicine 9, 452–458.

Abe T, Kearns CF & Sato Y. (2006). Muscle size and strength are increased following walk training with restricted venous blood flow from the leg muscle, Kaatsu-walk training. Journal of applied physiology 100, 1460–1466.

Abe T, Yasuda T, Midorikawa T, Sato Y, Kearns CF, Inoue K, Koizumi K & Ishii N. (2005). Skeletal muscle size and circulating IGF-1 are increased after two weeks of twice daily" KAATSU" resistance training. Int j KAATSU Training Res 1.

Adriano Pereira L, Freitas V, Arruda Moura F, Saldanha Aoki M, Loturco I & Yuzo Nakamura F. (2016) The Activity Profile of Young Tennis Athletes Playing on Clay and Hard Courts: Preliminary Data. Journal of human kinetics 50, 211–218.

Aughey RJ, Murphy KT, Clark SA, Garnham AP, Snow RJ, Cameron-Smith D, Hawley JA & McKenna MJ. (2007). Muscle Na+-K+-ATPase activity and isoform adaptations to intense interval exercise and training in well-trained athletes. Journal of applied physiology 103, 39–47.

Bangsbo J. (1994). The physiology of soccer-with special reference to intense intermittent exercise. Acta physiologica Scandinavica Supplementum 619, 1–155.

Birk JB & Wojtaszewski JF. (2006). Predominant alpha2/beta2/gamma3 AMPK activation during exercise in human skeletal muscle. The Journal of physiology 577, 1021–1032.

Bishop D, Jenkins DG & Mackinnon LT. (1998). The relationship between plasma lactate parameters, Wpeak and 1-h cycling performance in women. Medicine and science in sports and exercise 30, 1270–1275.

Bishop DJ. (2012). Fatigue during intermittent-sprint exercise. Clinical and experimental pharmacology & physiology 39, 836–841.

Bishop DJ & Girard O. (2013). Determinants of team-sport performance: implications for altitude training by team-sport athletes. British journal of sports medicine 47 Suppl 1, i17–21.

Bishop DJ, Granata C & Eynon N. (2014). Can we optimise the exercise training prescription to maximise improvements in mitochondria function and content? Biochimica et biophysica acta 1840, 1266–1275.

Brandt N, Gunnarsson TP, Hostrup M, Tybirk J, Nybo L, Pilegaard H & Bangsbo J. (2016). Impact of adrenaline and metabolic stress on exercise-induced intracellular signaling and PGC-1alpha mRNA response in human skeletal muscle. Physiological reports 4.

Briggs FN, Lee KF, Wechsler AW & Jones LR. (1992). Phospholamban expressed in slow-twitch and chronically stimulated fast-twitch muscles minimally affects calcium affinity of sarcoplasmic reticulum Ca(2+)-ATPase. The Journal of biological chemistry 267, 26056–26061.

Chen ZP, McConell GK, Michell BJ, Snow RJ, Canny BJ & Kemp BE. (2000). AMPK signaling in contracting human skeletal muscle: acetyl-CoA carboxylase and NO synthase phosphorylation. American journal of physiology Endocrinology and metabolism 279, E1202–1206.

Clanton TL. (2007). Hypoxia-induced reactive oxygen species formation in skeletal muscle. Journal of applied physiology 102, 2379–2388.

Clausen T. (2013). Quantification of Na+,K+ pumps and their transport rate in skeletal muscle: functional significance. The Journal of general physiology 142, 327–345.

Cohen J. (1988). Statistical power analysis for the behavioral sciences. 2nd edition.

Cook CJ, Kilduff LP & Beaven CM. (2014). Improving strength and power in trained athletes with 3 weeks of occlusion training. International journal of sports physiology and performance 9, 166–172.

Edge J, Eynon N, McKenna MJ, Goodman CA, Harris RC & Bishop DJ. (2013). Altering the rest interval during high-intensity interval training does not affect muscle or performance adaptations. Experimental physiology 98, 481–490.

Egan B, Carson BP, Garcia-Roves PM, Chibalin AV, Sarsfield FM, Barron N, McCaffrey N, Moyna NM, Zierath JR & O'Gorman DJ. (2010). Exercise intensity-dependent regulation of peroxisome proliferator-activated receptor coactivator-1 mRNA abundance is associated with differential activation of upstream signalling kinases in human skeletal muscle. The Journal of physiology 588, 1779–1790.

Eiken O & Bjurstedt H. (1987). Dynamic exercise in man as influenced by experimental restriction of blood flow in the working muscles. Acta physiologica Scandinavica 131, 339–345.

Ernst E & Resch KL. (1995). Concept of true and perceived placebo effects. Bmj 311, 551–553.

Fujita T, Brechue WF, Kurita K, Sato Y & Abe T. (2008). Increased muscle volume and strength following six days of low-intensity resistance training with restricted muscle blood flow. Int j KAATSU Training Res, 1–8.

Greenhaff PL, Soderlund K, Ren JM & Hultman E. (1993). Energy metabolism in single human muscle fibres during intermittent contraction with occluded circulation. The Journal of physiology 460, 443–453.

Gundermann DM, Fry CS, Dickinson JM, Walker DK, Timmerman KL, Drummond MJ, Volpi E & Rasmussen BB. (2012). Reactive hyperemia is not responsible for stimulating muscle protein synthesis following blood flow restriction exercise. Journal of applied physiology 112, 1520–1528.

Harris RC, Hultman E & Nordesjo LO. (1974). Glycogen, glycolytic intermediates and high-energy phosphates determined in biopsy samples of musculus quadriceps femoris of man at rest. Methods and variance of values. Scandinavian journal of clinical and laboratory investigation 33, 109–120.

Hashimoto T, Hussien R, Oommen S, Gohil K & Brooks GA. (2007). Lactate sensitive transcription factor network in L6 cells: activation of MCT1 and mitochondrial biogenesis. FASEB journal : official publication of the Federation of American Societies for Experimental Biology 21, 2602–2612.

Horiuchi M & Okita K. (2012). Blood flow restricted exercise and vascular function. International journal of vascular medicine 2012, 543218.

Hutchinson DS & Bengtsson T. (2006). AMP-activated protein kinase activation by adrenoceptors in L6 skeletal muscle cells: mediation by alpha1-adrenoceptors causing glucose uptake. Diabetes 55, 682–690.

Irrcher I, Ljubicic V & Hood DA. (2009). Interactions between ROS and AMP kinase activity in the regulation of PGC-1alpha transcription in skeletal muscle cells. American journal of physiology Cell physiology 296, C116–123.

Irrcher I, Ljubicic V, Kirwan AF & Hood DA. (2008). AMP-activated protein kinase-regulated activation of the PGC-1alpha promoter in skeletal muscle cells. PloS one 3, e3614.

Juel C, Hostrup M & Bangsbo J. (2015). The effect of exercise and beta2-adrenergic stimulation on glutathionylation and function of the Na,K-ATPase in human skeletal muscle. Physiological reports 3.

Kasai K, Yamashita T, Yamaguchi A, Yoshiya K, Kawakita A, Tanaka H, Sugimoto H & Tohyama M. (2003). Induction of mRNAs and proteins for Na/K ATPase alpha1 and beta1 subunits following hypoxia/reoxygenation in astrocytes. Brain research Molecular brain research 110, 38–44.

Kemp TJ, Causton HC & Clerk A. (2003). Changes in gene expression induced by H(2)O(2) in cardiac myocytes. Biochemical and biophysical research communications 307, 416–421.

Kourie JI. (1998). Interaction of reactive oxygen species with ion transport mechanisms. The American journal of physiology 275, C1–24.

Kristensen DE, Albers PH, Prats C, Baba O, Birk JB & Wojtaszewski JF. (2015). Human muscle fibre type-specific regulation of AMPK and downstream targets by exercise. The Journal of physiology.

Kubota A, Sakuraba K, Koh S, Ogura Y & Tamura Y. (2011). Blood flow restriction by low compressive force prevents disuse muscular weakness. Journal of science and medicine in sport / Sports Medicine Australia 14, 95–99.

Liu CC, Karimi Galougahi K, Weisbrod RM, Hansen T, Ravaie R, Nunez A, Liu YB, Fry N, Garcia A, Hamilton EJ, Sweadner KJ, Cohen RA & Figtree GA. (2013). Oxidative inhibition of the vascular Na+-K+ pump via NADPH oxidase-dependent beta1-subunit glutathionylation: implications for angiotensin II-induced vascular dysfunction. Free radical biology & medicine 65, 563–572.

Livak KJ & Schmittgen TD. (2001). Analysis of relative gene expression data using real-time quantitative PCR and the 2(-Delta Delta C(T)) Method. Methods (San Diego, Calif) 25, 402–408.

McKenna MJ. (1992). The roles of ionic processes in muscular fatigue during intense exercise. Sports medicine 13, 134–145.

McKenna MJ, Bangsbo J & Renaud JM. (2008). Muscle K+, Na+, and Cl disturbances and Na+-K+ pump inactivation: implications for fatigue. Journal of applied physiology 104, 288–295.

Murphy KT, Medved I, Brown MJ, Cameron-Smith D & McKenna MJ. (2008). Antioxidant treatment with N-acetylcysteine regulates mammalian skeletal muscle Na+-K+-ATPase alpha gene expression during repeated contractions. Experimental physiology 93, 1239–1248.

Murphy KT, Snow RJ, Petersen AC, Murphy RM, Mollica J, Lee JS, Garnham AP, Aughey RJ, Leppik JA, Medved I, Cameron-Smith D & McKenna MJ. (2004). Intense exercise up-regulates Na+,K+-ATPase isoform mRNA, but not protein expression in human skeletal muscle. The Journal of physiology 556, 507–519.

Murphy RM. (2011). Enhanced technique to measure proteins in single segments of human skeletal muscle fibers: fiber-type dependence of AMPK-alpha1 and -beta1. Journal of applied physiology 110, 820–825.

Nielsen JJ, Mohr M, Klarskov C, Kristensen M, Krustrup P, Juel C & Bangsbo J. (2004). Effects of high-intensity intermittent training on potassium kinetics and performance in human skeletal muscle. The Journal of physiology 554, 857–870.

Nilsson EC, Long YC, Martinsson S, Glund S, Garcia-Roves P, Svensson LT, Andersson L, Zierath JR & Mahlapuu M. (2006). Opposite transcriptional regulation in skeletal muscle of AMP-activated protein kinase gamma3 R225Q transgenic versus knock-out mice. The Journal of biological chemistry 281, 7244–7252.

Nordsborg N, Thomassen M, Lundby C, Pilegaard H & Bangsbo J. (2005). Contraction-induced increases in Na+-K+-ATPase mRNA levels in human skeletal muscle are not amplified by activation of additional muscle mass. American journal of physiology Regulatory, integrative and comparative physiology 289, R84–91.

Nordsborg NB, Kusuhara K, Hellsten Y, Lyngby S, Lundby C, Madsen K & Pilegaard H. (2010a). Contraction-induced changes in skeletal muscle Na(+), K(+) pump mRNA expression - importance of exercise intensity and Ca(2+)-mediated signalling. Acta physiologica 198, 487–498.

Nordsborg NB, Lundby C, Leick L & Pilegaard H. (2010b). Relative workload determines exercise-induced increases in PGC-1alpha mRNA. Medicine and science in sports and exercise 42, 1477–1484.

Norrbom J, Sundberg CJ, Ameln H, Kraus WE, Jansson E & Gustafsson T. (2004). PGC-1alpha mRNA expression is influenced by metabolic perturbation in exercising human skeletal muscle. Journal of applied physiology 96, 189–194.

Ojuka EO, Jones TE, Han DH, Chen M & Holloszy JO. (2003). Raising Ca2+ in L6 myotubes mimics effects of exercise on mitochondrial biogenesis in muscle. FASEB journal : official publication of the Federation of American Societies for Experimental Biology 17, 675–681.

Orlowski J & Lingrel JB. (1988). Tissue-specific and developmental regulation of rat Na,K-ATPase catalytic alpha isoform and beta subunit mRNAs. The Journal of biological chemistry 263, 10436–10442.

Perry CG, Lally J, Holloway GP, Heigenhauser GJ, Bonen A & Spriet LL. (2010). Repeated transient mRNA bursts precede increases in transcriptional and mitochondrial proteins during training in human skeletal muscle. The Journal of physiology 588, 4795–4810.

Povoas SC, Seabra AF, Ascensao AA, Magalhaes J, Soares JM & Rebelo AN. (2012). Physical and physiological demands of elite team handball. Journal of strength and conditioning research/ National Strength & Conditioning Association 26, 3365–3375.

Radzyukevich TL, Neumann JC, Rindler TN, Oshiro N, Goldhamer DJ, Lingrel JB & Heiny JA. (2013). Tissue-specific role of the Na,K-ATPase alpha2 isozyme in skeletal muscle. The Journal of biological chemistry 288, 1226–1237.

Raedschelders K, Ansley DM & Chen DD. (2012). The cellular and molecular origin of reactive oxygen species generation during myocardial ischemia and reperfusion. Pharmacology & therapeutics 133, 230–255.

Ristow M, Zarse K, Oberbach A, Kloting N, Birringer M, Kiehntopf M, Stumvoll M, Kahn CR & Bluher M. (2009). Antioxidants prevent health-promoting effects of physical exercise in humans. Proceedings of the National Academy of Sciences of the United States of America 106, 8665–8670.

Rose AJ, Kiens B & Richter EA. (2006). Ca(2+)–calmodulin-dependent protein kinase expression and signalling in skeletal muscle during exercise. The Journal of physiology 574, 889–903.

Ruas JL, White JP, Rao RR, Kleiner S, Brannan KT, Harrison BC, Greene NP, Wu J, Estall JL, Irving BA, Lanza IR, Rasbach KA, Okutsu M, Nair KS, Yan Z, Leinwand LA & Spiegelman BM. (2012). A PGC-1alpha isoform induced by resistance training regulates skeletal muscle hypertrophy. Cell 151, 1319–1331.

Silva E & Soares-da-Silva P. (2007). Reactive oxygen species and the regulation of renal Na+-K+-ATPase in opossum kidney cells. American journal of physiology Regulatory, integrative and comparative physiology 293, R1764–1770.

Slezak J, Tribulova N, Pristacova J, Uhrik B, Thomas T, Khaper N, Kaul N & Singal PK. (1995). Hydrogen peroxide changes in ischemic and reperfused heart. Cytochemistry and biochemical and X-ray microanalysis. The American journal of pathology 147, 772–781.

Slivka DR, Heesch MW, Dumke CL, Cuddy JS, Hailes WS & Ruby BC. (2014). Human skeletal muscle mRNAResponse to a single hypoxic exercise bout. Wilderness & environmental medicine 25, 462–465.

Stoner JD, Clanton TL, Aune SE & Angelos MG. (2007). O2 delivery and redox state are determinants of compartment-specific reactive O2 species in myocardial reperfusion. American journal of physiology Heart and circulatory physiology 292, H109–116.

Sundberg CJ. (1994). Exercise and training during graded leg ischaemia in healthy man with special reference to effects on skeletal muscle. Acta physiologica Scandinavica Supplementum 615, 1–50.

Sundberg CJ & Kaijser L. (1992). Effects of graded restriction of perfusion on circulation and metabolism in the working leg; quantification of a human ischaemia-model. Acta physiologica Scandinavica 146, 1–9.

Tavi P, Allen DG, Niemela P, Vuolteenaho O, Weckstrom M & Westerblad H. (2003). Calmodulin kinase modulates Ca2+ release in mouse skeletal muscle. The Journal of physiology 551, 5–12.

Taylor CW, Ingham SA, Hunt JE, Martin NR, Pringle JS & Ferguson RA. (2016). Exercise duration-matched interval and continuous sprint cycling induce similar increases in AMPK phosphorylation, PGC-1alpha and VEGF mRNA expression in trained individuals. European journal of applied physiology 116, 1445–1454.

Thomas C, Sirvent P, Perrey S, Raynaud E & Mercier J. (2004). Relationships between maximal muscle oxidative capacity and blood lactate removal after supramaximal exercise and fatigue indexes in humans. Journal of applied physiology 97, 2132–2138.

Van Beekvelt MC, Colier WN, Wevers RA & Van Engelen BG. (2001). Performance of near-infrared spectroscopy in measuring local O(2) consumption and blood flow in skeletal muscle. Journal of applied physiology 90, 511–519.

Vandesompele J, De Preter K, Pattyn F, Poppe B, Van Roy N, De Paepe A & Speleman F. (2002). Accurate normalization of real-time quantitative RT-PCR data by geometric averaging of multiple internal control genes. Genome biology 3, Research0034.

Varley MC, Gabbett T & Aughey RJ. (2013). Activity profiles of professional soccer, rugby league and Australian football match play. Journal of sports sciences.

Wang G, Kawakami K & Gick G. (2007). Regulation of Na,K-ATPase alpha1 subunit gene transcription in response to low K(+): role of CRE/ATF-and GC box-binding proteins. Journal of cellular physiology 213, 167–176.

Wendt CH, Sharma R, Bair R, Towle H & Ingbar DH. (1998). Oxidant effects on epithelial Na,K-ATPase gene expression and promoter function. Environmental health perspectives 106 Suppl 5, 1213–1217.

Wu H, Kanatous SB, Thurmond FA, Gallardo T, Isotani E, Bassel-Duby R & Williams RS. (2002).Regulation of mitochondrial biogenesis in skeletal muscle by CaMK. Science (New York, NY) 296, 349–352.

Wyckelsma VL, McKenna MJ, Serpiello FR, Lamboley CR, Aughey RJ, Stepto NK, Bishop DJ & Murphy RM. (2015). Single fiber expression and fiber-specific adaptability to short-term intense exercise training of Na+,K+-ATPase alpha and beta isoforms in human skeletal muscle. Journal of applied physiology, jap.00419.02014.

Zhang Y, Uguccioni G, Ljubicic V, Irrcher I, Iqbal S, Singh K, Ding S & Hood DA. (2014). Multiple signaling pathways regulate contractile activity-mediated PGC-1alpha gene expression and activity in skeletal muscle cells. Physiological reports 2.

Zhong RZ, Zhou DW, Tan CY, Tan ZL, Han XF, Zhou CS & Tang SX. (2011). Effect of tea catechins on regulation of antioxidant enzyme expression in H2O2-induced skeletal muscle cells of goat in vitro. Journal of agricultural and food chemistry 59, 11338–11343.

Zhuang Y, Wendt C & Gick G. (2000). Regulation of Na,K-ATPase beta 1 subunit gene transcription by low external potassium in cardiac myocytes. Role of Sp1 AND Sp3. The Journal of biological chemistry 275, 24173–24184.

